## Supplementary Figures and Note for "Quantifying portable genetic effects and improving cross-ancestry genetic prediction with GWAS summary statistics"

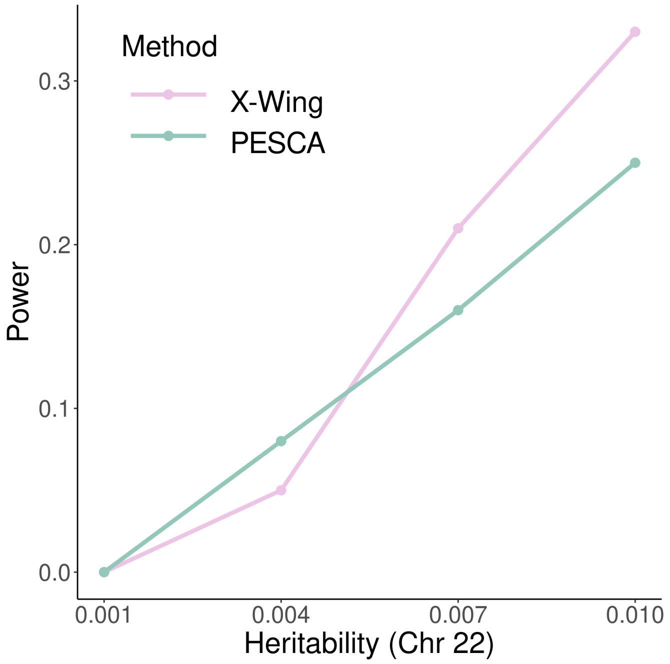

**Supplementary Figure 1. Statistical power in simulations under LDAK model in which SNP heritability is dependent on LD and MAF.**

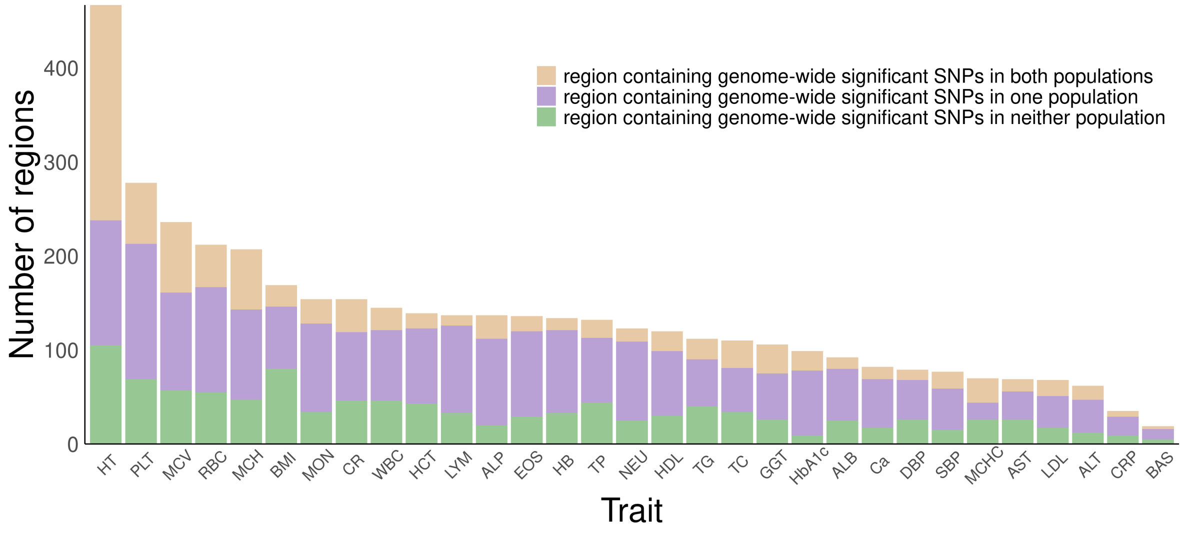

**Supplementary Figure 2. Number of regions with significant local genetic correlations between Europeans and East Asians in 31 complex traits.** Three bars denote regions containing genome-wide significant SNPs in both populations (brown), in only one population (purple), and in neither population (green).

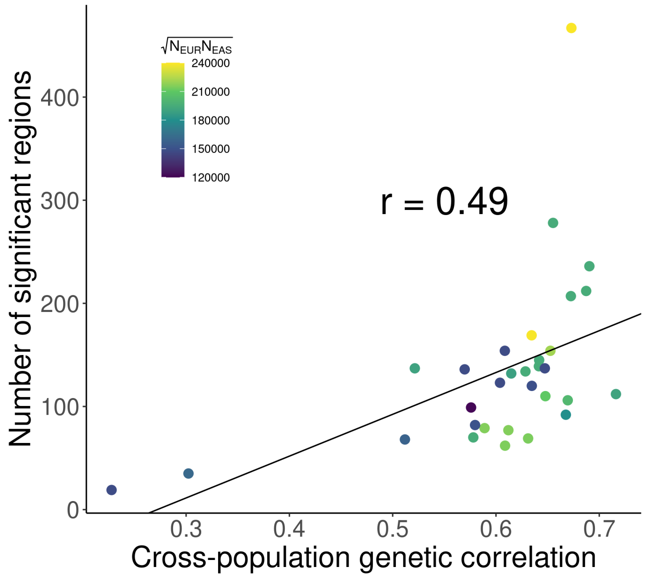

**Supplementary Figure 3. Number of X-Wing-identified regions is proportional to cross-population genetic correlation**. GWAS sample size are indicated by the color of each data point.

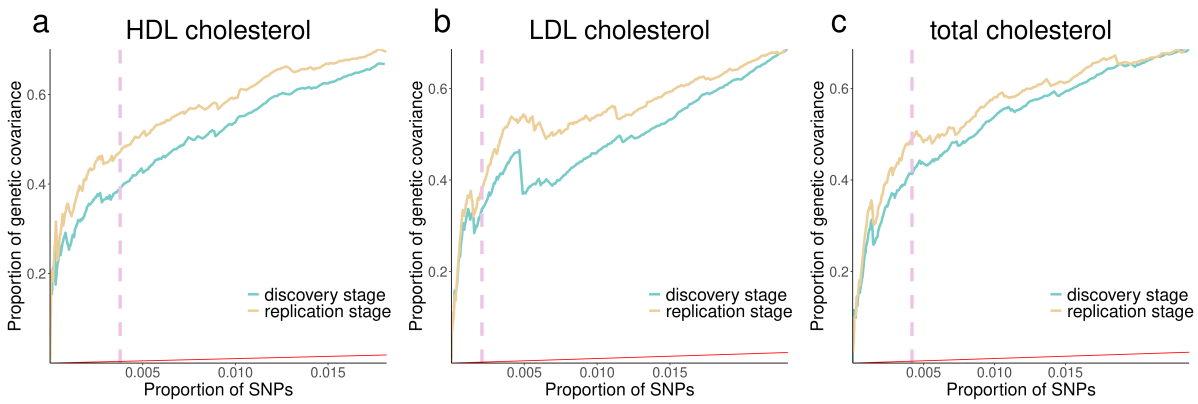

**Supplementary Figure 4. Cumulative proportion of genetic covariance explained by regions identified in the discovery stage for HDL cholesterol, LDL cholesterol, and total cholesterol**. Pink dashed line indicates FDR cutoff of 0.05. Red line indicates diagonal line of y=x.

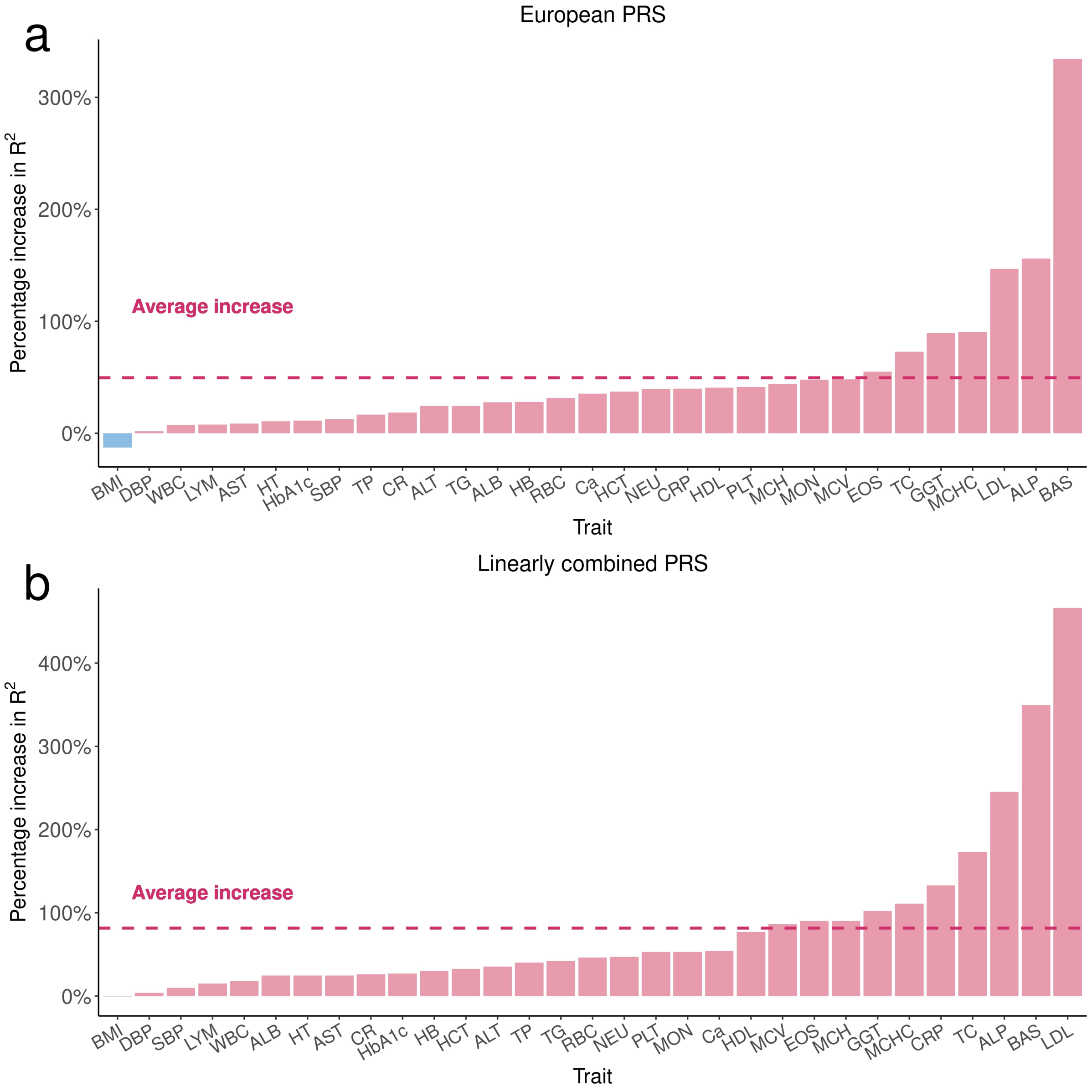

**Supplementary Figure 5. Comparison of the prediction accuracy between X-Wing and XPASS PRS for 31 traits in East Asian sample.** Panels **a** and **b** illustrate the percentage increase in $R^2$ of X-Wing European and linearly combined PRS over XPASS, respectively. The dashed line represents the average increase.

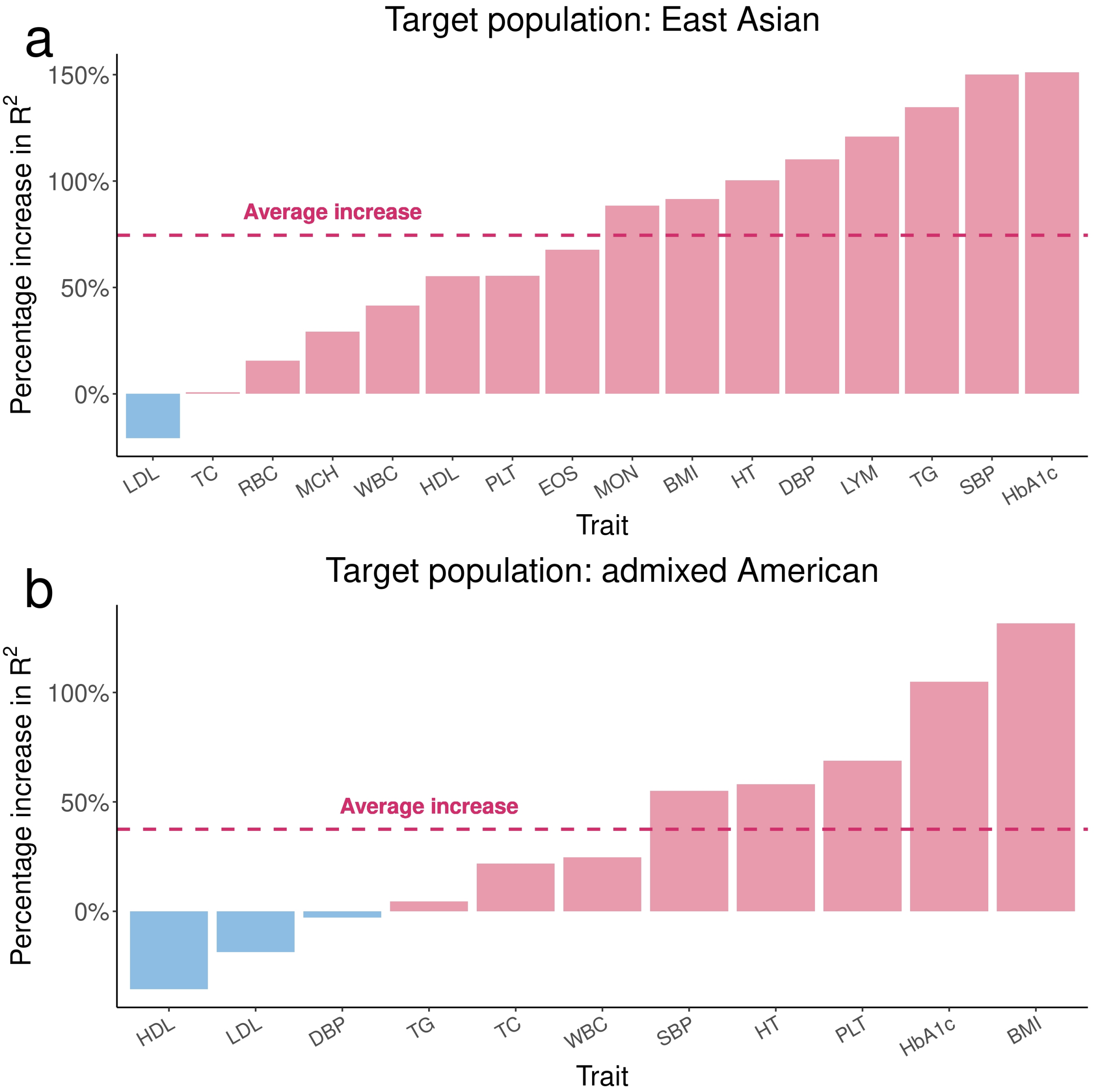

**Supplementary Figure 6. Comparison of the prediction accuracy between X-Wing and PolyFun-pred European PRS.** Panels **a** and **b** show the percentage increase in $R^2$ of X-Wing European PRS over PolyFun-pred for 16 traits in East Asians and 11 traits in admixed Americans, respectively.

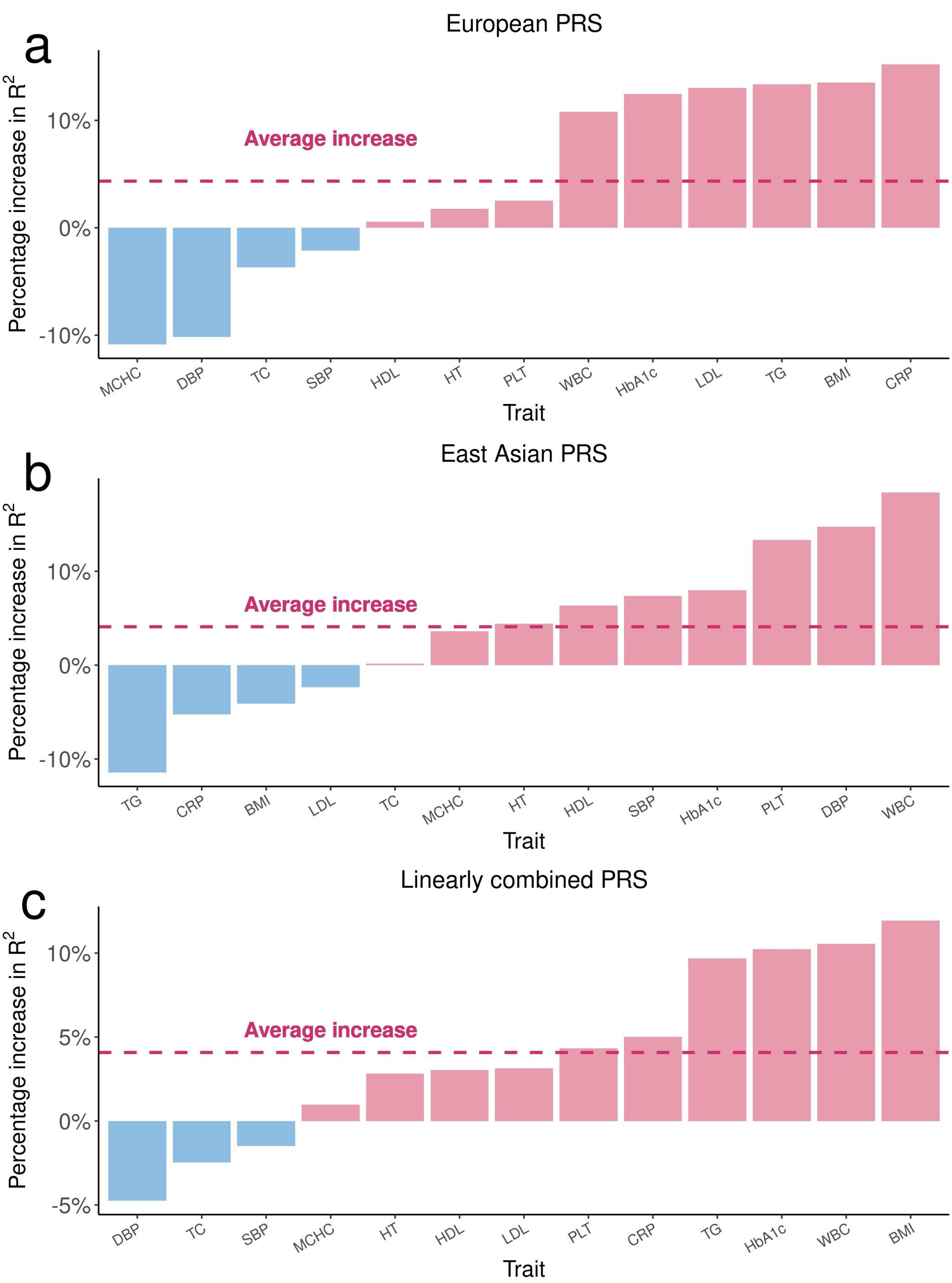

**Supplementary Figure 7. Comparison of the prediction accuracy between X-Wing and PRS-CSx PRS for 13 traits in admixed American sample.** Panels **a, b,** and **c** illustrate the percentage increase in $R^2$ of X-Wing European, East Asian, and linearly combined PRS over PRS-CSx. respectively. The dashed line represents the average increase.

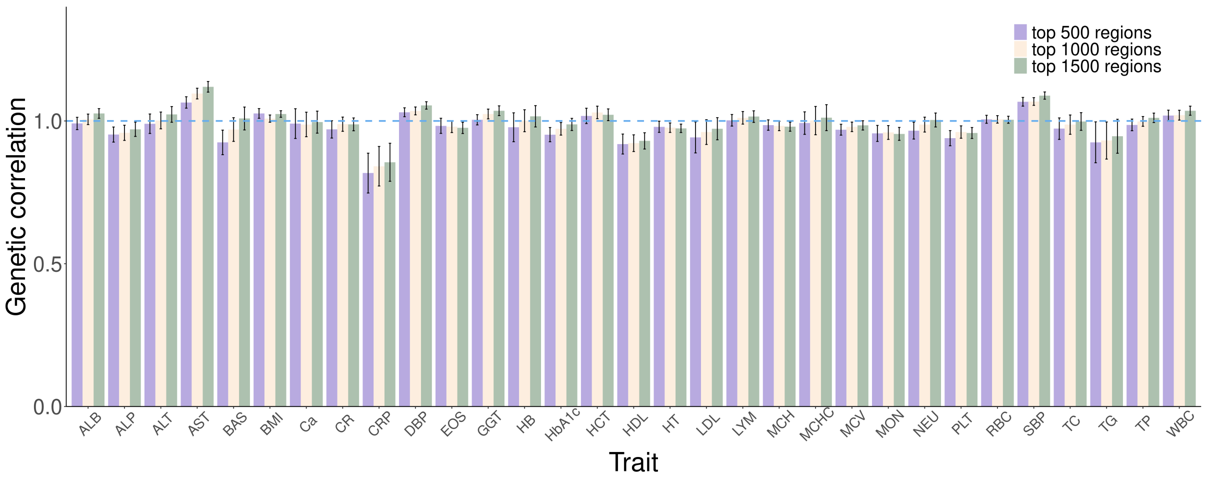

**Supplementary Figure 8. Bar plot shows the cross-population genetic correlation estimates and standard errors using varying numbers of top annotated regions.**

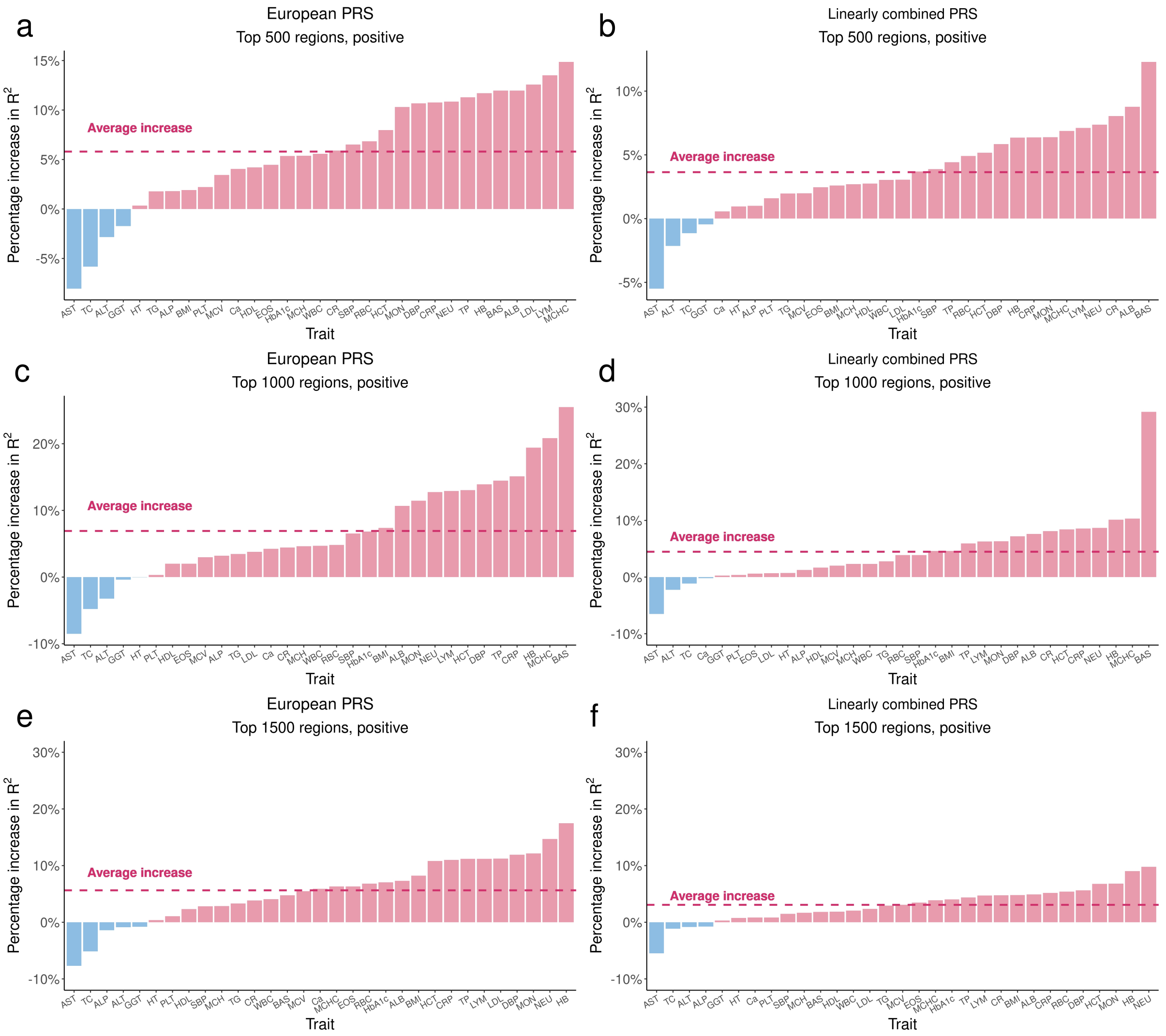

**Supplementary Figure 9. Comparison of the prediction accuracy between X-Wing and PRS-CSx PRS for 31 traits in East Asians with varying numbers of top regions.** Panel **a, c**, and **e** show the percentage increase in $R^2$ of X-Wing European PRS over PRS-CSx using 500, 1000, and 1500 positive regions as annotation. Panel **b, d**, and **f** are the results for linearly combined PRS. The dashed line represents the average increase.

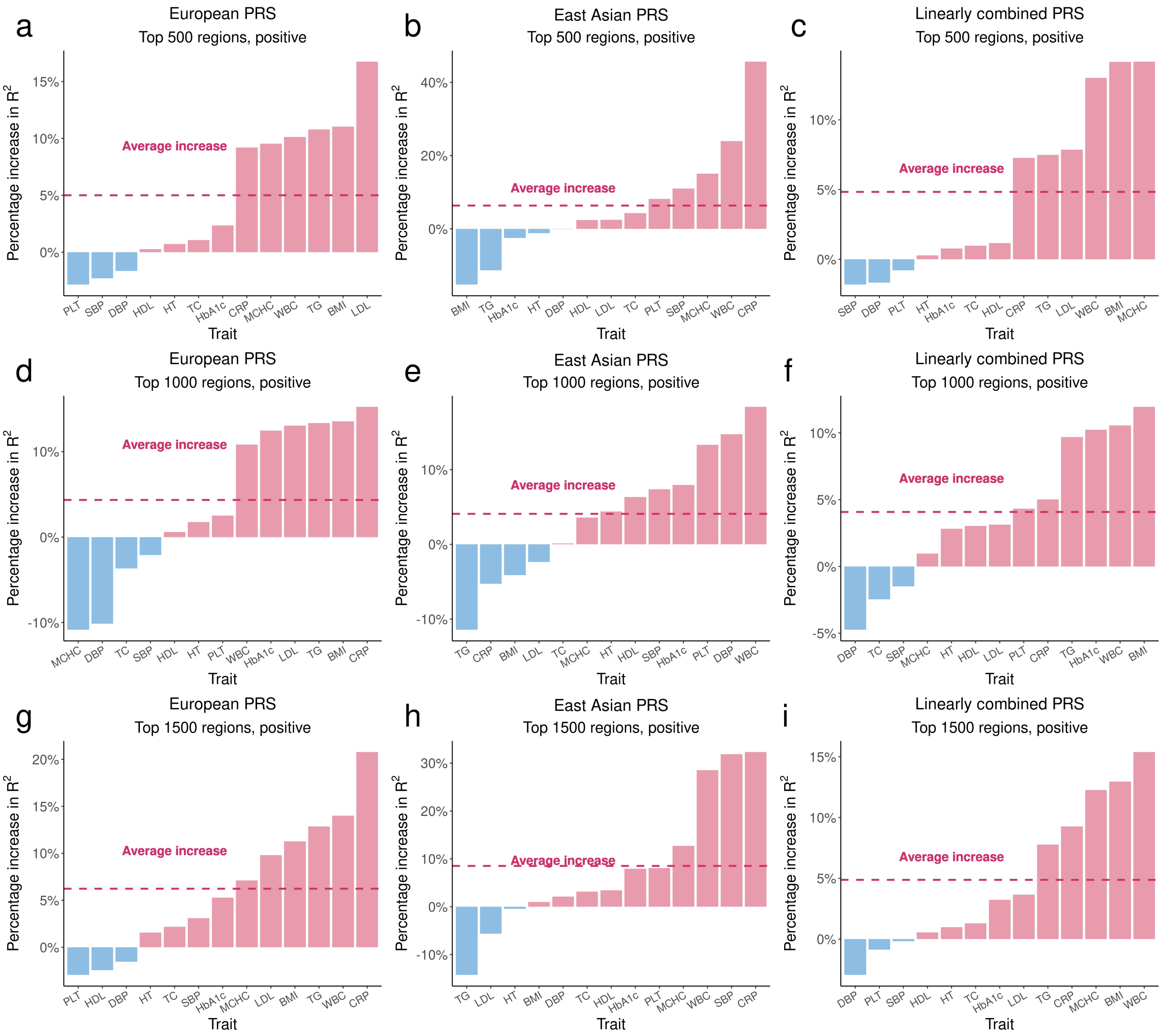

**Supplementary Figure 10. Comparison of the prediction accuracy between X-Wing and PRS-CSx PRS for 13 traits in admixed Americans with varying numbers of top regions.** Panel **a, d**, and **g** show the percentage increase in $R^2$ of European PRS from X-Wing over PRS-CSx using 500, 1000, and 1500 positive regions as annotation. Panel **b, e**, and **h** are the results for East Asian PRS. Panel **c, f**, and **i** represent the results for linearly combined PRS. The dashed line represents the average increase.

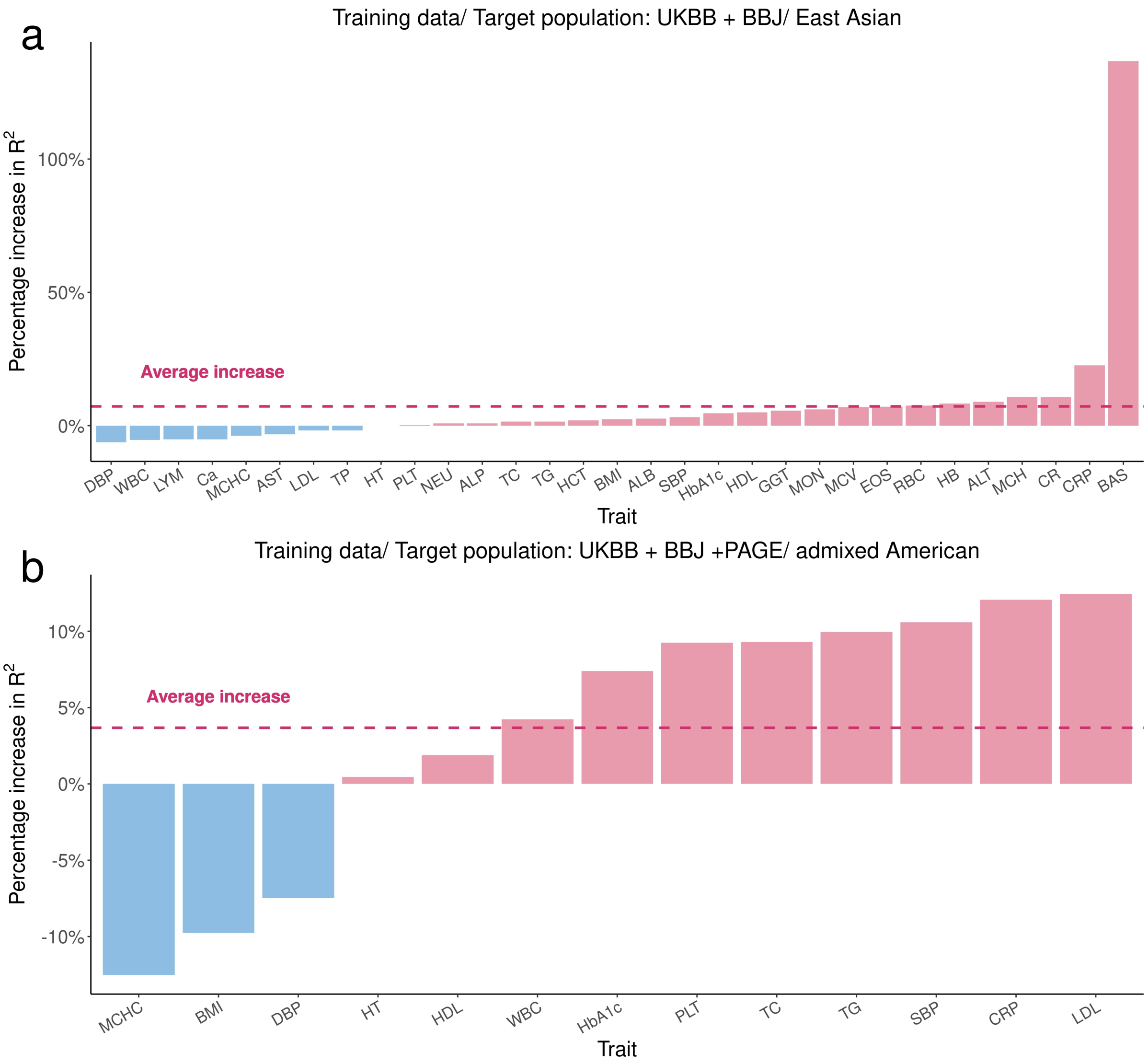

**Supplementary Figure 11. Comparison of the prediction accuracy between X-Wing and PRS-CSx PRS when using tuning parameter to select global shrinkage parameter.** The percentage increase in $R^2$ of linearly combined PRS from X-Wing over PRS-CSx is shown in **(a)** for 31 traits in East Asians and **(b)** for 13 traits in admixed Americans.

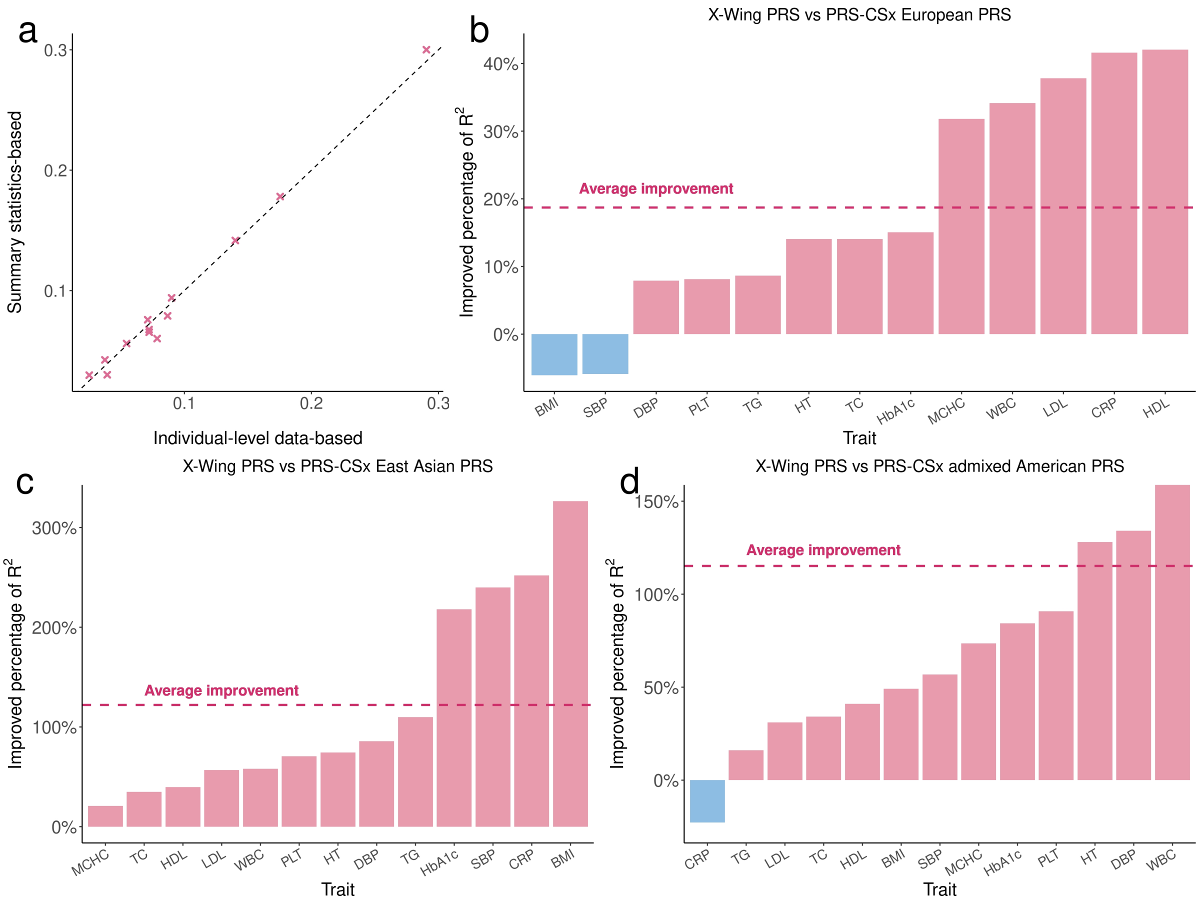

**Supplementary Figure 12. Performance of X-Wing in combining population-specific PRS using GWAS summary statistics for 13 traits in admixed Americans.** Panel **a** compares the $R^2$ for linearly combined PRS with mixing weights obtained using GWAS summary statistics and individual-level data. The X-axis represents the $R^2$ using weights estimated from individual-level data, while the Y-axis shows the $R^2$ using summary-statistics-based weights. The dashed line represents diagonal line of y=x. Panels **b, c**, and **d** show the percentage increase in $R^2$ of X-Wing PRS over PRS-CSx using only GWAS summary statistics, where the PRS-CSx PRS is calculated based on posterior mean effects of European, East Asian, and admixed American population, respectively. The dashed line represents the average increase.

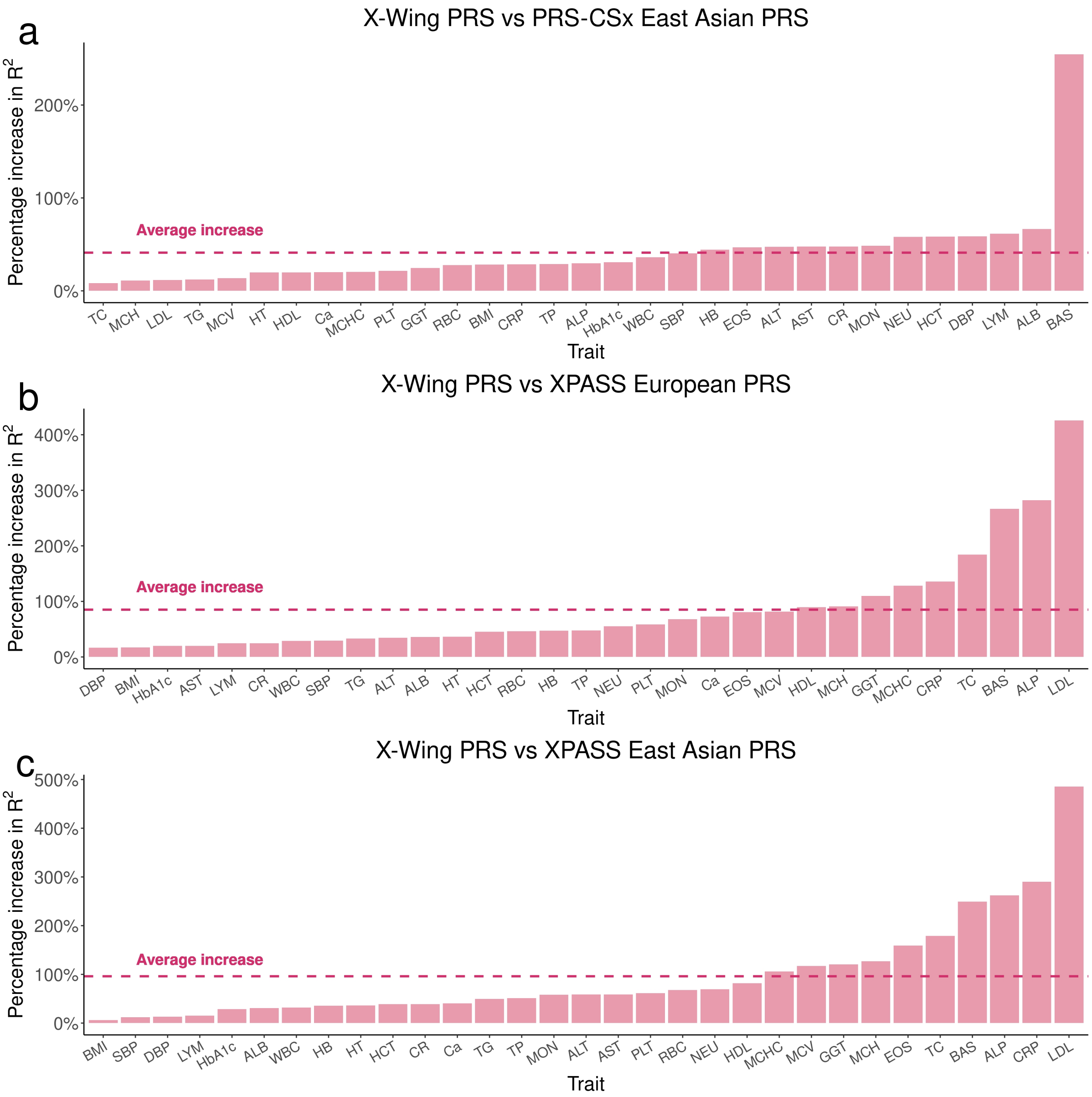

**Supplementary Figure 13. Comparison of performance of summary statistics-based linearly combined X-Wing PRS with other methods for 31 traits in East Asians.** It shows the percentage increase in $R^2$ of X-Wing PRS over PRS using **(a)** PRS-CSx East Asian **(b)** XPASS European **(c)** XPASS East Asian posterior mean effects**.**

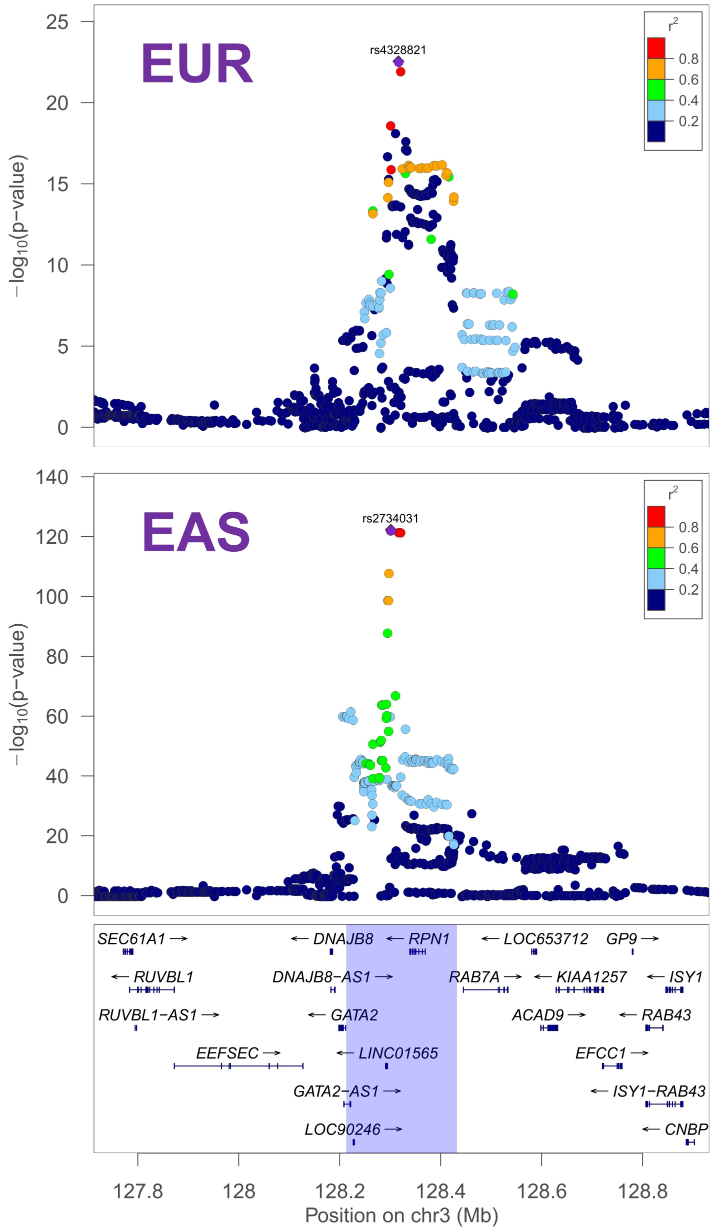

**Supplementary Figure 14. Locuszoom plot of a locus on chromosome 3 between Europeans and East Asians for basophil count**. The significant region is highlighted in blue. SNPs in this region achieve genome-wide significance in both populations (European lead SNP rs4328821, p = 2.82e-23; East Asian lead SNP rs2734031, p = 5.10e-123).

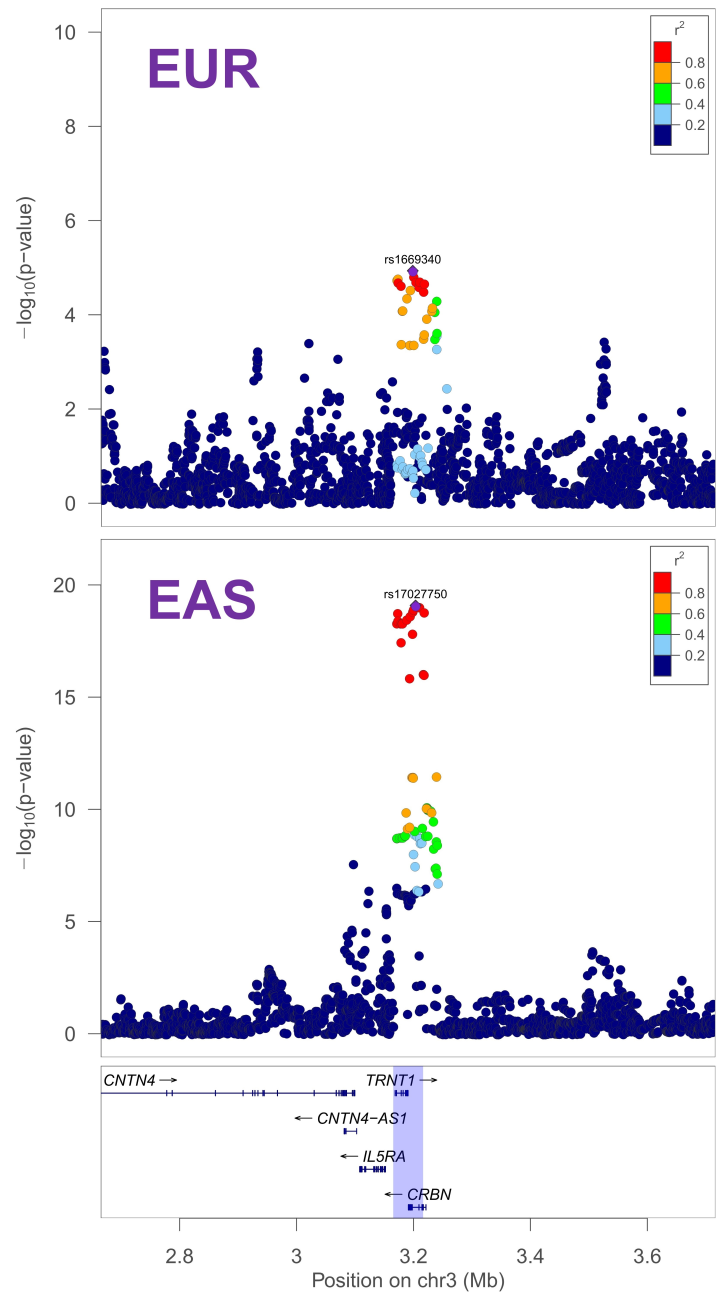

**Supplementary Figure 15. Locuszoom plot of a locus on chromosome 3 between Europeans and East Asians for basophil count.** The significant region is highlighted in blue. SNPs in this locus achieve genome-wide significance in East Asians (lead SNP rs17027750, p = 8.24e-20), but not in Europeans (lead SNP rs1669340, p = 1.16e-05).

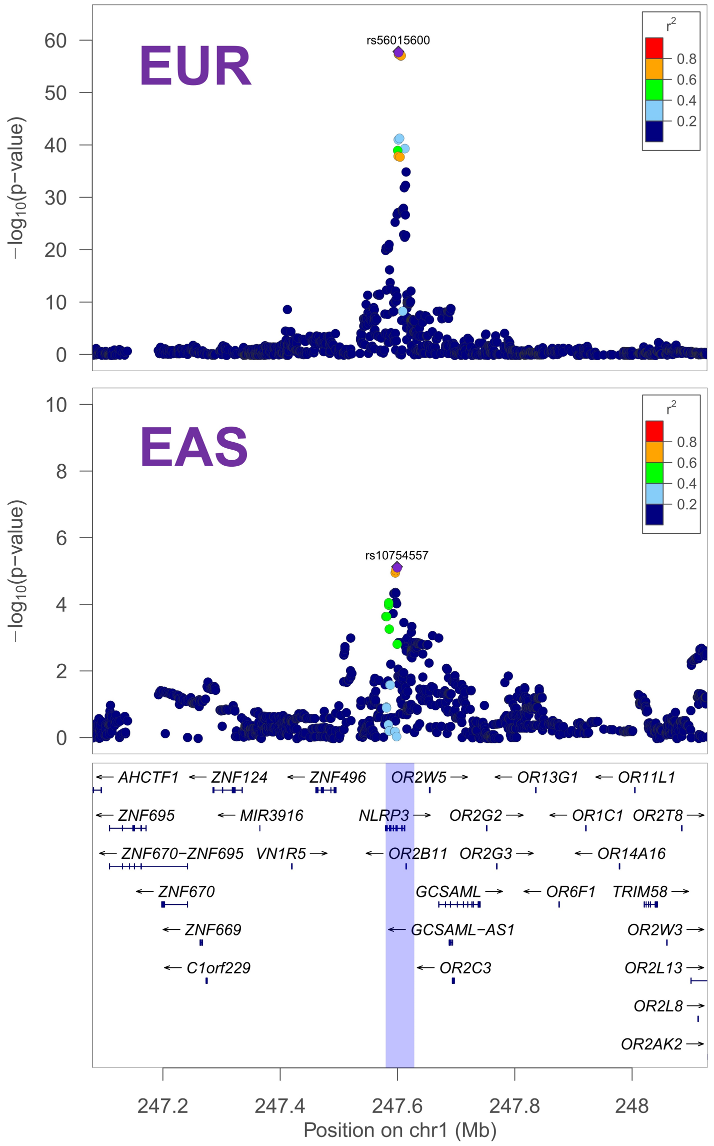

**Supplementary Figure 16. Locuszoom plot of a locus on chromosome 1 between Europeans and East Asians for C-reactive protein.** The significant region is highlighted in blue. SNPs in this locus achieve genome-wide significance in Europeans (lead SNP rs56015600, p = 1.48e-58), but not in East Asians (lead SNP rs10754557, p = 7.45e-6).

**Supplementary Note to**
**"Quantifying portable genetic effects and improving cross-ancestry genetic prediction with GWAS summary statistics"**

May 26, 2022

**4 Implementation of other methods 16**

### 1 Identification of genomic regions showing local genetic correlations between two ancestral populations

We have recently developed LOGODetect, which can precisely identify local, genetically-correlated regions between two traits in single population. Here we extend LOGODetect to detect genomic regions enriched for cross-population local genetic correlations.

Suppose the standardized traits $\boldsymbol{Y}_1$ and $\boldsymbol{Y}_2$ in two populations follow the linear models with random effects:

$$\boldsymbol{Y}_k = \boldsymbol{X}_k \boldsymbol{\beta}_k + \boldsymbol{\epsilon}_k, k = 1, 2,$$

where $\boldsymbol{X}_k$ is a $N_k \times M$ standardized genotype matrix; $\boldsymbol{\beta}_k$ is a $M$-dimensional vector of genetic effect sizes; $\boldsymbol{\epsilon}_k$ are non-genetic effects. We assume cross-population genetic covariance is localized in small regions $R_1, \ldots, R_r$, i.e. the joint genetic effect sizes follow the multivariate normal distribution:

$$\begin{bmatrix} \boldsymbol{\beta}_1 \\ \boldsymbol{\beta}_2 \end{bmatrix} \sim \mathcal{N}\left( \begin{bmatrix} \mathbf{0} \\ \mathbf{0} \end{bmatrix}, \begin{bmatrix} \frac{h_1^2}{M}\boldsymbol{I} & \frac{\rho_g}{K}\tilde{\boldsymbol{I}} \\ \frac{\rho_g}{K}\tilde{\boldsymbol{I}} & \frac{h_2^2}{M}\boldsymbol{I} \end{bmatrix} \right),$$

where $h_1^2$ and $h_2^2$ denote heritability in two populations; $\rho_g$ is the cross-population genetic covariance; $\tilde{\boldsymbol{I}}$ is a diagonal matrix where $\tilde{\boldsymbol{I}}[i,i] = 1$ if and only if $i \in \cup_{j=1}^r R_j$; $K = \sum_{j=1}^r |R_j|$, which is the number of SNPs with correlated genetic effect. We assume non-genetic effects $\boldsymbol{\epsilon}_k$ are independent across two populations.

Our goal is to use scan statistic to identify small regions enriched for local genetic correlation between two populations. Therefore, we design the numerator in the scan statistic as $\sum_{i \in R} z_{1i} z_{2i}$, the inner product of z-scores in a region across two populations, which quantifies the concordant association pattern of SNP effect sizes. To normalize the effects of LD in two populations, we use $\mathrm{Sd}\left[\sum_{i \in R} z_{1i} z_{2i}\right]$ under the null hypothesis as the denominator of scan statistic. Under the null hypothesis that cross-population genetic correlation is zero, the joint z-scores for two populations follow the multivariate normal distribution as

$$\begin{bmatrix} \boldsymbol{z}_1 \\ \boldsymbol{z}_2 \end{bmatrix} \sim \mathcal{N}\left( \begin{bmatrix} \mathbf{0} \\ \mathbf{0} \end{bmatrix}, \begin{bmatrix} \frac{\frac{h_1^2}{M}\boldsymbol{X}_1^T\boldsymbol{X}_1\boldsymbol{X}_1^T\boldsymbol{X}_1 + (1-h_1^2)\boldsymbol{X}_1^T\boldsymbol{X}_1}{n_1} & \mathbf{0} \\ \mathbf{0} & \frac{\frac{h_2^2}{M}\boldsymbol{X}_2^T\boldsymbol{X}_2\boldsymbol{X}_2^T\boldsymbol{X}_2 + (1-h_2^2)\boldsymbol{X}_2^T\boldsymbol{X}_2}{n_2} \end{bmatrix} \right).$$

Since individual genotype data is hardly accessible due to privacy issues in practice, we use LD matrices (denoted by $\boldsymbol{V}_k$) estimated from reference panels (e.g. the European and East Asian individuals from 1000 Genome Project) to approximate the sample LD matrices $\frac{\boldsymbol{X}_k^T\boldsymbol{X}_k}{N_k}$. Further, $\frac{\boldsymbol{X}_k^T\boldsymbol{X}_k\boldsymbol{X}_k^T\boldsymbol{X}_k}{N_k^2}$ can be unbiasedly estimated as $\widetilde{\boldsymbol{V}_k^2} = \frac{N_k^{(ref)}-1}{N_k^{(ref)}-2}\boldsymbol{V}_k^2 - \frac{M}{N_k^{(ref)}-2}\boldsymbol{V}_k$, where $N_k^{(ref)}$ is the sample size of the reference panel for population $k$. Using this approximation, we have

$$\begin{bmatrix} \boldsymbol{z}_1 \\ \boldsymbol{z}_2 \end{bmatrix} \sim \mathcal{N}\left( \begin{bmatrix} \mathbf{0} \\ \mathbf{0} \end{bmatrix}, \begin{bmatrix} \frac{N_1 h_1^2}{M} \widetilde{\boldsymbol{V}_1^2} + \left(1 - h_1^2\right) \boldsymbol{V}_1 & \mathbf{0} \\ \mathbf{0} & \frac{N_2 h_2^2}{M} \widetilde{\boldsymbol{V}_2^2} + \left(1 - h_2^2\right) \boldsymbol{V}_2 \end{bmatrix} \right).$$

We use XPASS (Cai et al., 2021) to estimate heritability for two populations $h_1^2$ and $h_2^2$. Let $\boldsymbol{\Sigma}_k = \frac{N_k h_k^2}{M} \widetilde{\boldsymbol{V}_k^2} + \left(1 - h_k^2\right) \boldsymbol{V}_k$. Denote $\boldsymbol{z}_{kR}$ as the sub-vector of $\boldsymbol{z}_k$ indexed by $R$, and $\boldsymbol{\Sigma}_{k,RR}$ as the sub-matrix of $\boldsymbol{\Sigma}_k$ whose rows and columns are both indexed by $R$. Then for a given region $R$, the joint z-scores in $R$ follows the multivariate normal distribution

$$\begin{bmatrix} \boldsymbol{z}_{1R} \\ \boldsymbol{z}_{2R} \end{bmatrix} \sim \mathcal{N}\left( \begin{bmatrix} \mathbf{0} \\ \mathbf{0} \end{bmatrix}, \begin{bmatrix} \boldsymbol{\Sigma}_{1,RR} & \mathbf{0} \\ \mathbf{0} & \boldsymbol{\Sigma}_{2,RR} \end{bmatrix} \right).$$

Following bilinear form theory (Searle and Gruber, 2016), we can show that $\mathrm{Var}\left[\sum_{i \in R} z_{1i} z_{2i}\right] = \mathrm{Tr}\left[\boldsymbol{\Sigma}_{1,RR} \boldsymbol{\Sigma}_{2,RR}\right]$, where $\mathrm{Tr}[\boldsymbol{A}]$ denotes the trace of matrix $\boldsymbol{A}$. However, computing value of $\mathrm{Tr}\left[\boldsymbol{\Sigma}_{1,RR} \boldsymbol{\Sigma}_{2,RR}\right]$ for all possible regions $R$ is computationally expensive. Therefore, we use $\sum_{i \in R} \boldsymbol{\Sigma}_{1,ii} * \boldsymbol{\Sigma}_{2,ii}$ to replace $\mathrm{Tr}\left[\boldsymbol{\Sigma}_{1,RR} \boldsymbol{\Sigma}_{2,RR}\right]$. We add tuning parameter $\theta$ to the power term of $\sum_{i \in R} \boldsymbol{\Sigma}_{1,ii} * \boldsymbol{\Sigma}_{2,ii}$ to accommodate the approximation bias and control the penalty strength of LD effects.

We select the best tuning parameter $\theta$ from a candidate set {0.5, 0.55, 0.6, 0.65, 0.7, 0.75} such that the identified regions for that given $\theta$ achieve the highest proportion of genetic covariance. The cross-population genetic covariance for aggregated regions can be estimated using XPASS (Cai et al., 2021).

We compute the cross-population LD matrix by taking the maximum of population-specific LD matrix in an element-wise fashion, as previously suggested (Shi et al., 2020). We use ldetect (Berisa and Pickrell, 2016) to divide the genome into 185 LD blocks (average size of 15MB) that are approximately independent in both populations based on the cross-population LD matrix. We first apply LOGODetect to identify regions with family-wise error rate cutoff of 0.05 in each LD block separately. Then we collect all the candidate regions identified across different LD blocks and control FDR level of 0.05 using the Benjamini-Hochberg procedure.

### 2 An annotation-dependent Bayesian horseshoe regression model for PRS

#### 2.1 Incorporating local genetic correlation annotation in PRS

##### 2.1.1 Model

Consider an additive genetic model:

$$\boldsymbol{Y}_k = \boldsymbol{X}_k \boldsymbol{\beta}_k + \boldsymbol{\epsilon}_k, \quad \boldsymbol{\epsilon}_k \sim \mathcal{MVN}\left(\mathbf{0}, \sigma_k^2 \boldsymbol{I}_k\right), \quad p\left(\sigma_k^2\right) \propto \sigma_k^{-2}, \quad k = 1, 2, \cdots, K,$$

where $\boldsymbol{\beta}_k$ is a $M$-dimensional vector of SNP effect sizes in population $k, \boldsymbol{\epsilon}_k$ is a vector of error terms with variance $\sigma_k^2$, to which we assign a non-informative Jeffreys prior. $\mathcal{MVN}$ denotes multivariate normal distribution, and $\boldsymbol{I}_k$ is an identity matrix.

Consider an annotation with $A$ category, we assign an annotation-dependent horseshoe prior to $\beta_{jk}$:

$$\beta_{jk} \sim \mathcal{N}\left(0, \frac{\sigma_k^2}{N_k} \phi \psi_j \lambda_{f(j),k}\right), \ j = 1, \ldots, M, \ k = 1, \ldots, K.$$

Here, $\beta_{jk}$ denotes the effect of SNP $j$ in population $k, \phi$ is the global shrinkage parameter shared across all $M$ SNPs, $\psi_j$ represents the local shrinkage parameters for SNP $j$, $\lambda_{f(j),k}$ denotes the annotation-dependent shrinkage parameter for SNP $j$ in population $k, f : j \rightarrow a \in \{1, \ldots A\}$ is a function that maps the j-th SNP to its corresponding category $a$ in the annotation.

To perform the full Bayesian model fitting, we assign the half-Cauchy priors to the global, local, and annotation-dependent shrinkage parameters as follows:

$$\begin{aligned}
&\phi^{\frac{1}{2}} \sim C^{+}(0,1) \\
&\psi_j^{\frac{1}{2}} \sim C^{+}(0,1) \quad j = 1, \ldots, M \\
&\lambda_{a,k}^{\frac{1}{2}} \sim C^{+}(0,1) \quad a = 1, \ldots, A, k = 1, \ldots, K.
\end{aligned}$$

Using the half-Cauchy decomposition, we have

$$\begin{aligned}
\phi \mid v &\sim \mathcal{IG}\left(\frac{1}{2}, \frac{1}{v}\right), v \sim \mathcal{IG}\left(\frac{1}{2}, 1\right) \\
\psi_j \mid c_j &\sim \mathcal{IG}\left(\frac{1}{2}, \frac{1}{c_j}\right), c_j \sim \mathcal{IG}\left(\frac{1}{2}, 1\right) \quad j = 1, \ldots, M \\
\lambda_{a,k} \mid t_{a,k} &\sim \mathcal{IG}\left(\frac{1}{2}, \frac{1}{t_{a,k}}\right), t_{a,k} \sim \mathcal{IG}\left(\frac{1}{2}, 1\right) \quad a = 1, \ldots, A,
\end{aligned}$$

where $\mathcal{IG}$ denotes the inverse-gamma distribution.

##### 2.1.2 Gibbs sampler

Next, we derive the full conditional distribution of all parameters in the above model. For notation purpose, we rewrite the prior in matrix form:

$$\boldsymbol{\beta}_k \sim \mathcal{N}\left(0, \frac{\sigma_k^2}{N_k}\phi\boldsymbol{\Psi}\boldsymbol{\Lambda}_k\right),\ j = 1, \ldots, M,\ k = 1, \ldots, K,$$

where $\boldsymbol{\Psi} = \mathrm{diag}\,(\psi_1, \ldots, \psi_M)\,, \boldsymbol{\Lambda}_k = \mathrm{diag}\left(\lambda_{f(1),k}, \ldots, \lambda_{f(M),k}\right)$.

The Gibbs sampler involves the following steps in each Markov Chain Monte Carlo (MCMC) iteration:

- update $\boldsymbol{\beta}_k : \boldsymbol{\beta}_k \mid \cdot \sim \mathcal{MVN}(\mu_k, \boldsymbol{\Sigma}_k), \quad \mu_k = \frac{N_k}{\sigma_k^2}\boldsymbol{\Sigma}_k\hat{\boldsymbol{\beta}}_k, \quad \boldsymbol{\Sigma}_k = \frac{\sigma_k^2}{N_k}\left(\boldsymbol{D}_k + (\phi\boldsymbol{\Psi}\boldsymbol{\Lambda}_k)^{-1}\right)^{-1},$

  where $D_k$ is the LD-matrix for population $k$, $\hat{\boldsymbol{\beta}}_k$ is the marginal least squares estimates obtained from GWAS summary statistics. To avoid the numerical issue caused by colinearity between SNPs, we restrict $(\phi\boldsymbol{\Psi}\boldsymbol{\Lambda}_k)^{-1} \geq 1$.

- update $\sigma_k^2 : \sigma_k^2 \mid \cdot \sim \mathcal{IG}\left(\frac{N_k+M_k}{2}, \frac{N_k}{2}\left[1 - 2\boldsymbol{\beta}_k^T\hat{\boldsymbol{\beta}}_k + \boldsymbol{\beta}_k^T\left(\boldsymbol{D}_k + \phi\boldsymbol{\Psi}\boldsymbol{\Lambda}_k\right)\boldsymbol{\beta}_k\right]\right),$

  where $M_k$ is number of SNPs in population $k$.

- update $\phi : \phi \mid \cdot \sim \mathcal{IG}\left(\frac{\sum_{k=1}^K M_k+2}{2}, \sum_{k=1}^K \frac{\boldsymbol{\beta}_k^T(\boldsymbol{\Psi}\boldsymbol{\Lambda}_k)^{-1}\boldsymbol{\beta}_k N_k}{2\sigma_k^2} + \frac{1}{\nu}\right)$

- update $\nu : \nu \mid \cdot \sim \mathcal{IG}\left(1, \frac{1}{\phi} + 1\right)$

- update $\psi_j : \psi_j \mid \cdot \sim \mathcal{IG}\left(\frac{k_j+1}{2}, \sum_{k=1}^K \frac{\beta_{jk}^2 N_k}{2\sigma_k^2\phi\lambda_{f(j),k}} + \frac{1}{c_j}\right)$

  where $k_j = 1$ if SNP $j$ exists only in one population and $r$ if it exists in $r$ populations included.

- update $c_j : c_j \mid \cdot \sim \mathcal{IG}\left(1, \frac{1}{\psi_j} + 1\right)$

- update $\lambda_{a,k} : \lambda_{a,k} \mid \cdot \sim \mathcal{IG}\left(\frac{s_q+1}{2}, \frac{1}{2\phi}\sum_{j \in l(a,k)} \frac{\beta_{jk}^2 N_k}{\psi_j\sigma_k^2} + \frac{1}{t_{a,k}}\right)$

  where $s_a$ is the number of predictors in category $a$, $l(a,k) = \{j \in \{1, \ldots, M\} : \lambda_{f(j),k} = \lambda_{a,k}\}$ is the set of SNP predictors that belongs to category $a$.

- update $t_{a,k} \mid \cdot \sim \mathcal{IG}\left(1, \frac{1}{\lambda_{a,k}} + 1\right)$

##### 2.1.3 Example of annotation-dependent shrinkage based on local genetic correlation annotation

Here, we provide an example of the annotation-dependent shrinkage $\lambda_{f(j),k}$ based on local genetic correlation annotation. WLOG, we assume that we have $K = 3$ populations in total and population $1$ as the target population.

Given the local genetic correlation annotation $\Omega_2$ and $\Omega_3$, the $\lambda_{f(j),k}$ in the Full Bayesian model fitting process is specified as:

| Annotation-dependent shrinkage $\lambda_{f(j),k}$ | **Population 1 (target)** | Population 2 | Population 3 |
|---|---|---|---|
| Posterior effects for population 1 | $1$ for all $j$ | $1$ for all $j$ | $1$ for all $j$ |
| Posterior effects for population 2 | $\lambda_{1,1}$ for all $j$ | $\lambda_{1,2}$ if SNP $j$ is not annotated by $\Omega_2$<br>$\lambda_{2,2}$ if SNP $j$ is annotated by $\Omega_2$ | $\lambda_{1,3}$ for all $j$ |
| Posterior effects for population 3 | $\lambda_{1,1}$ for all $j$ | $\lambda_{1,2}$ for all $j$ | $\lambda_{1,3}$ if SNP $j$ is not annotated by $\Omega_3$<br>$\lambda_{2,3}$ if SNP $j$ is annotated by $\Omega_3$ |

Here, $k$-th row represents the specification of the annotation-dependent shrinkage parameter $\lambda_{f(j),k'}$ for $k^{'} = 1, 2, 3$ when obtaining the posterior effects for $k$-th population.

##### 2.1.4 Model-tuning strategy

Instead of assigning a prior for $\phi$, we select the global shrinkage parameter $\phi$ from a grid of value $\{10^{-6}, 10^{-4}, 10^{-2}, 1\}$ with respect to the largest $R^2$ in the validation set. The detailed algorithm is listed below:

---

**Algorithm 1:** Model-tuning X-Wing

---

**Input:** GWAS summary statistics and population-matached LD reference panel from population 1 to $K$, target sample genotype.

**Output:** X-Wing PRS.

1 We perform local genetic correlation analysis between population 1 and population $k$ ($k = 2, \dots K$) to identify top $s$ regions with positive local genetic correlation. We denote the set of regions as $\Omega_k$.

2 For each $\phi \in \{10^{-6}, 10^{-4}, 10^{-2}, 1\}$, we fit our PRS model with annotation-dependent shrinkage specified below: when estimating the posterior SNP effects for the non-target population $k$ that $k \neq 1$, we used $\lambda_{f(j),k} = \lambda_{1,k}$ if SNP $j$ is not annotated by $\Omega_k$, $\lambda_{f(j),k} = \lambda_{2,k}$ if SNP $j$ is annotated by $\Omega_k$, and $\lambda_{f(j),k'} = 1$ for $k^{'} = 1, 2, \dots, k-1, k+1, \dots, K$. When estimating the posterior SNP effects for target population, we used $\lambda_{f(j),k} = 1$ for all $j = 1, 2, \dots M, k = 1, \dots K$.

3 For each $\phi \in \{10^{-6}, 10^{-4}, 10^{-2}, 1\}$, based on the posterior mean effects of population $k$ obtained in step2, we calculate population-specific score $\boldsymbol{PRS}_{k,\phi}$. A common practice to combine these population-specific scores is to fit a regression model using the same phenotype $\boldsymbol{Y}_1^{(v)}$ and K population-specific PRS in an independent validation dataset from the target population:

$$\boldsymbol{Y}_1^{(v)} \sim w_{1,\phi}\boldsymbol{PRS}_{1,\phi}^{(v)} + w_{2,\phi}\boldsymbol{PRS}_{2,\phi}^{(v)} + \cdots + w_{K,\phi}\boldsymbol{PRS}_{K,\phi}^{(v)},$$

Instead of fitting a regression in independent samples, we employ a novel strategy to obtain the least squares estimates of regression weights (i.e. $\widehat{w}_{1,\hat{\phi}}, \dots \widehat{w}_{K,\hat{\phi}}$) using GWAS summary statistics. We introduce this approach in the section 3.3.

4 The final X-Wing PRS is then calculated as:

$$\boldsymbol{PRS}_{LC} = \sum_{k=1}^{K} \widehat{w}_{k,\hat{\phi}}\boldsymbol{PRS}_{k,\hat{\phi}}$$

---

#### 2.2 Incorporating multiple annotations

##### 2.2.1 Model

We generalized our model to incorporate $T$ annotations with the annotation-dependent shrinkage prior to $\beta_{jk}$:

$$\beta_{jk} \sim \mathcal{N}\left(0, \frac{\sigma_k^2}{N_k}\phi_k\psi_j \prod_{t=1}^{T} \lambda_{f(j,t),k}\right),\ j = 1, \ldots, M,\ k = 1, \ldots, K.$$

Here, $\beta_{jk}$ denotes the effect of SNP $j$ in population $k, \phi_k$ is the global shrinkage parameter shared across all SNPs for population $k, \psi_j$ represents the local shrinkage parameters for SNP $j$ and are shared across population, $\lambda_{f(j,t),k}$ is the annotation-dependent shrinkage parameters for SNP $j$ in population $k$ for t-th annoration, $f : (j,t) \to a_t \in \{1, \ldots, \ldots A_t\}$ is a function that maps the $j$-th SNP to its corresponding category $a_t$ in the $t$-th annotation.

To perform the full Bayesian model fitting, we assign the half-Cauchy priors to the global, local, and annotation-dependent shrinkage parameters as follows:

$$\begin{aligned}
&\phi_k^{\frac{1}{2}} \sim C^+(0,1) \quad k = 1, \ldots, K \\
&\psi_j^{\frac{1}{2}} \sim C^+(0,1) \quad j = 1, \ldots, M \\
&\lambda_{a_t,k}^{\frac{1}{2}} \sim C^+(0,1) \quad a_t = 1, \ldots, A_t, t = 1, \ldots, T, k = 1, \ldots, K
\end{aligned}$$

Using the half-Cauchy decomposition, we have

$$\begin{aligned}
\phi_k \mid v &\sim \mathcal{IG}\left(\frac{1}{2}, \frac{1}{v_k}\right), v_k \sim \mathcal{IG}\left(\frac{1}{2}, 1\right) \\
\psi_j \mid c_j &\sim \mathcal{IG}\left(\frac{1}{2}, \frac{1}{c_j}\right), c_j \sim \mathcal{IG}\left(\frac{1}{2}, 1\right) \quad j = 1, \ldots, M \\
\lambda_{a_t,k} \mid t_{a_t,k} &\sim \mathcal{IG}\left(\frac{1}{2}, \frac{1}{t_{a_t,k}}\right), t_{a_t,k} \sim \mathcal{IG}\left(\frac{1}{2}, 1\right) \quad a_t = 1, \ldots, A_t, k = 1, \ldots, K,
\end{aligned}$$

where $\mathcal{IG}$ denotes the inverse-gamma distribution.

##### 2.2.2 Gibbs sampler

Next, we derive the full conditional distribution of all parameters in the above model. For notation purpose, we rewrite the prior in matrix form:

$$\boldsymbol{\beta}_k \sim \mathcal{N}\left(0, \frac{\sigma_k^2}{N_k}\phi_k \boldsymbol{\Psi}\boldsymbol{\Lambda}_{k1} \cdots \boldsymbol{\Lambda}_{kT}\right),\ j = 1, \ldots, M,\ k = 1, \ldots, K,$$

where $\boldsymbol{\Psi} = \text{diag}(\psi_1, \ldots, \psi_M), \boldsymbol{\Lambda}_{kT} = \text{diag}\left(\lambda_{f(1,t),k}, \ldots, \lambda_{f(M,t),k}\right), t = 1, \ldots, T.$

The Gibbs sampler then involves the following steps in each MCMC iteration:

- update $\boldsymbol{\beta}_k : \boldsymbol{\beta}_k \mid \cdot \sim \mathcal{MVN}(\mu_k, \boldsymbol{\Sigma}_k), \quad \mu_k = \frac{N_k}{\sigma_k^2}\boldsymbol{\Sigma}_k\hat{\boldsymbol{\beta}}_k, \quad \boldsymbol{\Sigma}_k = \frac{\sigma_k^2}{N_k}\left(\boldsymbol{D}_k + (\phi_k\boldsymbol{\Psi}\boldsymbol{\Lambda}_{k1}\cdots\boldsymbol{\Lambda}_{kT})^{-1}\right)^{-1},$

where $D_k$ is the LD-matrix for population $k$, $\hat{\boldsymbol{\beta}}_k$ is the marginal least squares estimates obtained from GWAS summary statistics. To avoid the numerical issue caused by colinearity between SNPs, we restrict $(\phi_k \boldsymbol{\Psi} \boldsymbol{\Lambda}_{k1} \cdots \boldsymbol{\Lambda}_{kT})^{-1} \geq 1$.

- update $\sigma_k^2 : \sigma_k^2 \mid \cdot \sim \mathcal{IG}\left(\frac{N_k+M_k}{2}, \frac{N_k}{2}\left[1 - 2\boldsymbol{\beta}_k^T \hat{\boldsymbol{\beta}}_k + \boldsymbol{\beta}_k^T \left(\boldsymbol{D}_k + \phi_k \boldsymbol{\Psi} \boldsymbol{\Lambda}_{k1} \cdots \boldsymbol{\Lambda}_{kT}\right) \boldsymbol{\beta}_k\right]\right)$
- update $\phi_k : \phi_k \mid \cdot \sim \mathcal{IG}\left(\frac{M_k+2}{2}, \frac{\boldsymbol{\beta}_k^T (\boldsymbol{\Psi} \boldsymbol{\Lambda}_{k1} \cdots \boldsymbol{\Lambda}_{kT})^{-1} \boldsymbol{\beta}_k N_k}{2\sigma_k^2} + \frac{1}{\nu}\right)$
- update $\nu_k : \nu_k \mid \cdot \sim \mathcal{IG}\left(1, \frac{1}{\phi_k} + 1\right)$
- update $\psi_j : \psi_j \mid \cdot \sim \mathcal{IG}\left(\frac{k_j+1}{2}, \sum_{k=1}^{K} \frac{\beta_{jk}^2 N_k}{2\sigma_k^2 \phi_k \prod_{t=1}^{T} \lambda_{f(j,t),k}} + \frac{1}{c_j}\right)$

  where $k_j = 1$ if SNP $j$ exists only in one population and $r$ if it exists in $r$ populations included.
- update $c_j : c_j \mid \cdot \sim \mathcal{IG}\left(1, \frac{1}{\psi_j} + 1\right)$
- update $\lambda_{a_t,k} : \lambda_{a_t,k} \mid \cdot \sim \mathcal{IG}\left(\frac{s_{a_t}+1}{2}, \frac{1}{2\phi_k} \sum_{j \in l(a_t,k)} \frac{\beta_{jk}^2 N_k}{\prod_{t \neq t} \lambda_{f(j,t),k} \psi_j \sigma_k^2} + \frac{1}{t_{a,k}}\right)$

  where $s_{a_t}$ is the number of predictors in category $a_t$, $l(a_t, k) = \{j \in \{1, \ldots, M\} : \lambda_{f(j,t),k} = \lambda_{a_t,k}\}$ is the set of predictors belonging to category $a_t$.
- update $t_{a_t,k} \mid \cdot \sim \mathcal{IG}\left(1, \frac{1}{\lambda_{a_t,k}} + 1\right)$

### 3 Combining multiple PRS with GWAS summary statistics

#### 3.1 Sufficient statistics for least squares estimator of linear combination weights

Consider the linear combination problem for $K$ centered population-specific PRS using the individual-level validation data $\left(\boldsymbol{X}_1^{(v)}, \boldsymbol{Y}_1^{(v)}\right)$ with sample size $N_1^{(v)}$:

$$\boldsymbol{Y}_1^{(v)} \sim \boldsymbol{PRS}^{(v)} \boldsymbol{w}.$$

Here, superscript $v$ highlights the fact that phenotypes and PRS in this regression exercise need to be obtained from a validation dataset that is different from any data used for GWAS and PRS training. $\boldsymbol{Y}_1^{(v)}$ is the phenotype vector and $\boldsymbol{PRS}^{(v)}$ is the $N_1^{(v)} \times K$ matrix of $K$ population-specific scores in this sample. Further, $\boldsymbol{PRS}^{(v)}$ can be denoted as $\boldsymbol{PRS}^{(v)} = \boldsymbol{X}_1^{(v)} \boldsymbol{b}$ where $\boldsymbol{X}_1^{(v)}$ is the $N_1^{(v)} \times M$ genotype matrix and $\boldsymbol{b}$ is the $M \times K$ matrix for SNP effects, $\boldsymbol{w} = [w_1, \ldots, w_K]^T$ is a $K$-dimensional linear combination weights vector.For simplicity, we assume $\boldsymbol{Y}_1^{(v)}$ is centered, $\boldsymbol{X}_1^{(v)}$ is standardized, and $\boldsymbol{b}$ quantifies standardized SNP effects.

Next, we showed the The least squares estimator for $\boldsymbol{w}$ is

$$\begin{aligned}\widehat{\boldsymbol{w}} &= \left[\boldsymbol{PRS}^{(v)T}\boldsymbol{PRS}^{(v)}\right]^{-1}\boldsymbol{PRS}^{(v)T}\boldsymbol{Y}_1^{(v)} \\ &= \left[\boldsymbol{b}^T\boldsymbol{X}_1^{(v)T}\boldsymbol{X}_1^{(v)}\boldsymbol{b}\right]^{-1}\boldsymbol{b}^T\boldsymbol{X}_1^{(v)T}\boldsymbol{Y}_1^{(v)}.\end{aligned}$$

This indicates that $\boldsymbol{b}, \boldsymbol{X}_1^{(v)T}\boldsymbol{X}_1^{(v)}$, and $\boldsymbol{X}_1^{(v)T}\boldsymbol{Y}_1^{(v)}$ are sufficient statistics for $\widehat{\boldsymbol{w}}$, where $\boldsymbol{b}$ is obtained from the PRS training procedure, $\boldsymbol{X}_1^{(v)T}\boldsymbol{X}_1^{(v)}$ is from in-sample LD matrix, and $\boldsymbol{X}_1^{(v)T}\boldsymbol{Y}_1^{(v)}$ can be obtained from the summary statistics of the validation sample. When the in-sample LD information is not available, we use LD matrix from the reference panel $\frac{\boldsymbol{X}^{(ref)T}\boldsymbol{X}^{(ref)}}{N^{(ref)}}$ as replacement. Then we have

$$\begin{aligned}\widehat{\boldsymbol{w}} &= \left[\boldsymbol{b}^T\boldsymbol{X}_1^{(v)T}\boldsymbol{X}_1^{(v)}\boldsymbol{b}\right]^{-1}\boldsymbol{b}^T\boldsymbol{X}_1^{(v)T}\boldsymbol{Y}_1^{(v)} \\ &\approx \left[\frac{N_1^{(v)}}{N^{(ref)}}\boldsymbol{b}^T\boldsymbol{X}^{(ref)T}\boldsymbol{X}^{(ref)}\boldsymbol{b}\right]^{-1}\boldsymbol{b}^T\boldsymbol{X}_1^{(v)T}\boldsymbol{Y}_1^{(v)} \\ &= \frac{N^{(ref)}}{N_1^{(v)}}\left[\boldsymbol{PRS}^{(ref)T}\boldsymbol{PRS}^{(ref)}\right]^{-\mathbf{1}}\boldsymbol{b}^T\boldsymbol{X}_1^{(v)T}\boldsymbol{Y}_1^{(v)},\end{aligned}$$

where $N^{(ref)}$ and $\boldsymbol{PRS}^{(ref)}$ denote the sample size and PRS matrix in the reference panel. Taken together, this shows that in order to obtain $\widehat{\boldsymbol{w}}$, we only need the LD reference and summary statistics from a validation sample.

#### 3.2 Derivation for subsampling GWAS summary statistics from training and validation sets

Consider the phenotype-genotype model:

$$Y_i = \boldsymbol{X_i}\boldsymbol{\beta} + \epsilon_i,$$

where $Y_i$ is the standardized phenotype with mean 0 and variance 1 for individual $i$, $\boldsymbol{X_i}$ is a $1 \times M$ standardized genotype matrix, and $\epsilon_i$ is the error term, $\boldsymbol{\beta}$ is a p-dimensional effect sizes vector. Note that the subscript $i$ in section 3.2 denotes the individual rather than the population.

Here, we consider $X_i$ and $Y_i$ as random and *i.i.d.* distributed (*i.e.*, $Y_1, \ldots, Y_N \overset{i.i.d}{\sim} Y_1 \in \mathbb{R}$, $\boldsymbol{X}_1, \ldots, \boldsymbol{X}_N \overset{i.i.d}{\sim} \boldsymbol{X}_1 \in \mathbb{R}^{M\times 1}$). We denote $\boldsymbol{Y} = (Y_1, \ldots, Y_N)^T$ as a $N$-dimensional phenotype vector and $\boldsymbol{X} = (\boldsymbol{X}_1^T, \ldots, \boldsymbol{X}_N^T)^T$ as a $N \times M$ standardized genotype matrix.

The standard approach the process for model validation technique involves first randomly sampling a subset of $N - N^{(v)}$ individuals from full sample $(\boldsymbol{X}, \boldsymbol{Y})$ as the training data $(\boldsymbol{X}^{(tr)}, \boldsymbol{Y}^{(tr)})$, and use the remaining $N^{(v)}$ individuals as the validation data $(\boldsymbol{X}^{(v)}, \boldsymbol{Y}^{(v)})$,

The GWAS sample size is large and hence by the central limit theorem, we have approximately

$$\boldsymbol{X}^T\boldsymbol{Y} \sim \mathcal{N}(N\mathbb{E}[\boldsymbol{X}_1^T Y_1], N\operatorname{Var}[\boldsymbol{X}_1^T Y_1])$$
$$\boldsymbol{X}^{(tr)T}\boldsymbol{Y}^{(tr)} \sim \mathcal{N}((N-N_1^{(v)})\mathbb{E}[\boldsymbol{X}_1^T Y_1], (N-N_1^{(v)})\operatorname{Var}[\boldsymbol{X}_1^T Y_1]).$$

The covariance between $\boldsymbol{X}^T\boldsymbol{Y}$ and $\boldsymbol{X}^{(tr)T}\boldsymbol{Y}^{(tr)}$ is

$$\begin{aligned}\operatorname{Cov}(\boldsymbol{X}^T\boldsymbol{Y}, \boldsymbol{X}^{(tr)T}\boldsymbol{Y}^{(tr)}) &= \operatorname{Cov}(\boldsymbol{X}^{(tr)T}\boldsymbol{Y}^{(tr)} + \boldsymbol{X}^{(v)T}\boldsymbol{Y}^{(v)}, \boldsymbol{X}^{(tr)T}\boldsymbol{Y}^{(tr)}) \\ &= \operatorname{Var}(\boldsymbol{X}^{(tr)T}\boldsymbol{Y}^{(tr)}) \\ &= (N-N^{(v)})\operatorname{Var}[\boldsymbol{X}_1^T Y_1].\end{aligned}$$

Here, we use the formula for the conditional distribution of two multivariate normal random vectors: if $\boldsymbol{A} \sim \mathcal{N}(\boldsymbol{\mu_A}, \boldsymbol{\Sigma_A})$, $\boldsymbol{B} \sim \mathcal{N}(\boldsymbol{\mu_B}, \boldsymbol{\Sigma_B})$, and $\operatorname{Cov}(\boldsymbol{A}, \boldsymbol{B}) = \boldsymbol{\Sigma_{AB}}$, we have the distribution of $\boldsymbol{A}|\boldsymbol{B}$ following a multivariate normal distribution with mean and covariance matrix.

$$\begin{aligned}\mathbb{E}[\boldsymbol{A} \mid \boldsymbol{B} = \boldsymbol{b}] &= \boldsymbol{\mu_A} + \boldsymbol{\Sigma_{AB}}\boldsymbol{\Sigma_B}^{-1}(\boldsymbol{b} - \boldsymbol{\mu_B}) \\ \operatorname{Var}[\boldsymbol{A} \mid \boldsymbol{B} = \boldsymbol{b}] &= \boldsymbol{\Sigma_A} - \boldsymbol{\Sigma_{AB}}\boldsymbol{\Sigma_B}^{-1}\boldsymbol{\Sigma_{AB}}.\end{aligned}$$

Thus, we have

$$\begin{aligned}\mathbb{E}[\boldsymbol{X}^{(tr)T}\boldsymbol{Y}^{(tr)} \mid \boldsymbol{X}^T\boldsymbol{Y} = \boldsymbol{x}^T\boldsymbol{y}] &= (N-N^{(v)})\mathbb{E}[\boldsymbol{X}_1^T Y_1] + \frac{N-N^{(v)}}{N}\left(\boldsymbol{x}^T\boldsymbol{y} - N\mathbb{E}[\boldsymbol{X}_1^T Y_1]\right) \\ \operatorname{Var}[\boldsymbol{X}^{(tr)T}\boldsymbol{Y}^{(tr)} \mid \boldsymbol{X}^T\boldsymbol{Y} = \boldsymbol{x}^T\boldsymbol{y}] &= (N-N^{(v)})\operatorname{Var}[\boldsymbol{X}_1^T Y_1] - \frac{N-N^{(v)}}{N}(N-N^{(v)})\operatorname{Var}[\boldsymbol{X}_1^T Y_1] \\ &= \frac{(N-N^{(v)})N^{(v)}}{N}\operatorname{Var}[\boldsymbol{X}_1^T Y_1].\end{aligned}$$

For the conditional expectation, we plug in the estimator $\boldsymbol{x}^T\boldsymbol{y}$ for $N\mathbb{E}[\boldsymbol{X}_1^T Y_1]$, the estimator for the conditional expectation is

$$\mathbb{E}[\boldsymbol{X}^{(tr)T}\widehat{\boldsymbol{Y}^{(tr)} \mid \boldsymbol{X}}^T\boldsymbol{Y} = \boldsymbol{x}^T\boldsymbol{y}] = \frac{N-N_1^{(v)}}{N}\boldsymbol{x}^T\boldsymbol{y}$$

For notation purpose, we define the conditional variance as

$$\operatorname{Var}[\boldsymbol{X}_1^T Y_1] := \boldsymbol{\Sigma} \in \mathbb{R}^{M\times M}$$

The diagonal term $\boldsymbol{\Sigma_{jj}}$ of $\boldsymbol{\Sigma}$ is

$$\begin{aligned}\boldsymbol{\Sigma_{jj}} &= \operatorname{Var}[X_{1j}Y_1] \\ &= \mathbb{E}[X_{1j}^2 Y_1^2] - \mathbb{E}[X_{1j}Y_1]^2 \\ &= \mathbb{E}[X_{1j}^2]\mathbb{E}[Y_1^2] + \operatorname{Cov}(X_{1j}^2, Y_1^2) - [\mathbb{E}[X_{1j}]\mathbb{E}[Y_1] + \operatorname{Cov}(X_{1j}, Y_1)]^2 \\ &\approx \mathbb{E}[X_{1j}^2] \\ &= 1\end{aligned}$$

The off-diagonal term $\boldsymbol{\Sigma_{jk}}$ of $\boldsymbol{\Sigma}, j \neq k$ is

$$
\begin{aligned}
\boldsymbol{\Sigma_{jk}} &= \mathrm{Cov}[X_{1j}Y_1, X_{1k}Y_1] \\
&= \mathbb{E}[X_{1j}X_{1k}Y_1^2] - \mathbb{E}[X_{1j}Y_1]\mathbb{E}[X_{1k}Y_1] \\
&= \mathbb{E}[X_{1j}X_{1k}]\mathbb{E}[Y_1^2] + \mathrm{Cov}(X_{1j}X_{1k}, Y_1^2) - [\mathbb{E}[X_{1j}]\mathbb{E}[Y_1] + \mathrm{Cov}(X_{1j}, Y_1)][\mathbb{E}[X_{1k}]\mathbb{E}[Y_1] + \mathrm{Cov}(X_{1k}, Y_1)] \\
&\approx \mathbb{E}[X_{1j}X_{1k}]
\end{aligned}
$$

It turns out that the $\boldsymbol{\Sigma}$ is exactly the LD matrix constructed using the standardized genotypes. Thus, we obtain the estimator for the conditional variance as

$$
\widehat{\mathrm{Var}[\boldsymbol{X}^{(tr)T}\boldsymbol{Y}^{(tr)} \mid \boldsymbol{X}^T\boldsymbol{Y} = \boldsymbol{x}^T\boldsymbol{y}]} = \frac{(N - N^{(v)})N^{(v)}}{N^{(v)}}\hat{\boldsymbol{\Sigma}},
$$

where $\hat{\boldsymbol{\Sigma}} = \frac{\boldsymbol{X}^{(ref)T}\boldsymbol{X}^{(ref)}}{N^{(ref)}}$ is obtained from the reference panel, $\boldsymbol{X}^{(ref)}$ is a $N^{(ref)} \times M$ standardized genotype matrix.

In conclusion, we have

$$
\boldsymbol{X}^{(tr)T}\boldsymbol{Y}^{(tr)} \mid \boldsymbol{X^T Y} = \boldsymbol{x^T y} \sim \mathcal{N}\left(\frac{(N - N^{(v)})}{N}\boldsymbol{x}^T\boldsymbol{y}, \frac{(N - N^{(v)})N^{(v)}}{N}\hat{\boldsymbol{\Sigma}}\right)
$$

Thus, we subsample the summary statistics for training set given full summary statistics $\boldsymbol{X^T Y}$ by

$$
\frac{\boldsymbol{X}^{(tr)T}\boldsymbol{Y}_1^{(tr)}}{N - N^{(v)}} \mid \boldsymbol{X^T Y} = \frac{\boldsymbol{X}^T\boldsymbol{Y}}{N} + \left(\frac{N^{(v)}}{N - N^{(v)}}\right)^{\frac{1}{2}} \frac{\boldsymbol{X}^{(ref)T}}{\sqrt{N^{(ref)}}}\boldsymbol{g}
$$

where $\boldsymbol{g}$ is a $N^{(ref)}$-dimensional vector with elements drawn from a standard normal distribution.

#### 3.3 Dealing with tuning parameters in the PRS model

If there are tuning parameters in the PRS model, we use the correlation $R$ between phenotype and linearly combined PRS in the validation set to select the optimal tuning parameter, as well as to estimate the linear combination weights.

Followed the notation above, suppose there are tuning parameters $\boldsymbol{\gamma}$ in the PRS model, consider the linear combination problem for $K$ centered PRS using the individual-level validation data:

$$
\boldsymbol{Y}_1^{(v)} \sim \boldsymbol{PRS}_{\gamma}^{(v)}\boldsymbol{w}_{\gamma}
$$

where $\boldsymbol{PRS}_{\gamma}^{(v)} = \boldsymbol{X}_1^{(v)}\boldsymbol{b}_{\gamma} = \left[\boldsymbol{PRS}_{1,\gamma}^{(v)}, \ldots \boldsymbol{PRS}_{k,\gamma}^{(v)}, \ldots, \boldsymbol{PRS}_{K,\gamma}^{(v)}\right]$ is a $N_1^{(v)} \times K$ centered PRS matrix with respect to to the tuning parameter $\boldsymbol{\gamma}$ in the PRS model, $\boldsymbol{w}_{\gamma}$ is a $K$-dimensional linear combination weights vector.

The least squares estimator for $\boldsymbol{w}_\gamma$ is

$$\hat{\boldsymbol{w}_\gamma} \approx \frac{N^{(ref)}}{N_1^{(v)}} \left[\boldsymbol{PRS}_\gamma^{(ref)T}\boldsymbol{PRS}_\gamma^{(ref)}\right]^{-1} \boldsymbol{b}_\gamma^{\boldsymbol{T}}\boldsymbol{X}_1^{(v)T}\boldsymbol{Y}_1^{(v)}$$

where $\boldsymbol{PRS}_\gamma^{(ref)} = \boldsymbol{X}^{(ref)}\boldsymbol{b}_\gamma$ is $N^{(ref)} \times K$ PRS matrix in the reference panel.

The correlation $R_\gamma$ between the linearly combined PRS and phenotype in the validation set with respect to to the estimated weights $\hat{\boldsymbol{w}_\gamma}$ is :

$$\begin{aligned} R_\gamma &= \frac{\hat{\boldsymbol{w}_\gamma}^T\boldsymbol{b}_\gamma^T\boldsymbol{X}_1^{(v)T}\boldsymbol{Y}_1^{(v)}/N_1^{(v)}}{\left(\hat{\boldsymbol{w}_\gamma}^T\boldsymbol{b}_\gamma^T\boldsymbol{X}_1^{(v)T}\boldsymbol{X}_1^{(v)}\boldsymbol{b}_\gamma\hat{\boldsymbol{w}_\gamma}/N_1^{(v)}\right)^{1/2}} \\ &\approx \frac{\hat{\boldsymbol{w}_\gamma}^T\boldsymbol{b}_\gamma^T\boldsymbol{X}_1^{(v)T}\boldsymbol{Y}_1^{(v)}/N_1^{(v)}}{\left(\hat{\boldsymbol{w}_\gamma}^T\boldsymbol{b}_\gamma^T\boldsymbol{X}_1^{(ref)T}\boldsymbol{X}_1^{(ref)}\boldsymbol{b}_\gamma\hat{\boldsymbol{w}_\gamma}/N^{(ref)}\right)^{1/2}} \\ &= \frac{\hat{\boldsymbol{w}_\gamma}^T\boldsymbol{b}_\gamma^T\boldsymbol{X}_1^{(v)T}\boldsymbol{Y}_1^{(v)}/N_1^{(v)}}{\left(\hat{\boldsymbol{w}_\gamma}^T\boldsymbol{PRS}_\gamma^{(ref)T}\boldsymbol{PRS}_\gamma^{(ref)}\hat{\boldsymbol{w}_\gamma}/N^{(ref)}\right)^{1/2}} \end{aligned}$$

Then we select the optimal tuning parameter $\hat{\gamma}$ as

$$\hat{\gamma} = \text{argmax}_\gamma R_\gamma,$$

and use the linear combination weights $\hat{\boldsymbol{w}}_{\hat{\gamma}}$ with respect to the optimal tuning parameter $\hat{\gamma}$ to linearly combine the PRS.

#### 3.4 Grid search to handle negative least squares estimates for mixing weights

In practice, the least squares estimates for linear combination weights of a particular PRS can be negative. It may decrease the prediction accuracy of the linearly combined PRS. Thus, we provide a grid search strategy to mimic the non-negative least squares.

We pre-specify a grid of positive value for the linear combination weights

$$\boldsymbol{w} \in \boldsymbol{W} = \{(w_1, \ldots, w_k, \ldots, w_K) \mid \sum_{k=1}^{K} w_k = 1, w_k \geq 0 \text{ for all } k = 1, \ldots, K\}.$$

Then, we use the formula in the above section to calculate the correlation between the linearly combined PRS and phenotype in the validation set. The linear combination weights with respect to to largest correlation, $\hat{\boldsymbol{w}}_{grid} = \text{argmax}_{\boldsymbol{w} \in \boldsymbol{W}} R_{\boldsymbol{w}}$, will be used to linearly combine the PRS.

#### 3.5 Summary statistics-based ridge regression to combine multiple PRS

When linearly combined many PRS with multicollinearity problems, the least squares of the linear combination weights may be sub-optimal. A remedy for multicollinearity is ridge regression.

We first describe a individual-level data-based ridge regression. Consider the linear combination problem for $K$ centered population-specific PRS using the individual-level validation data $\left(\boldsymbol{X}_1^{(v)}, \boldsymbol{Y}_1^{(v)}\right)$ with sample size $N_1^{(v)}$ :

$$\boldsymbol{Y}_1^{(v)} \sim \boldsymbol{PRS}^{(v)}\boldsymbol{w}.$$

Here, $\boldsymbol{Y}_1^{(v)}$ is the phenotype vector and $\boldsymbol{PRS}^{(v)}$ is the $N_1^{(v)} \times K$ matrix of $K$ population-specific scores in this sample. Further, $\boldsymbol{PRS}^{(v)}$ can be denoted as $\boldsymbol{PRS}^{(v)} = \boldsymbol{X}_1^{(v)}\boldsymbol{b}$ where $\boldsymbol{X}_1^{(v)}$ is the $N_1^{(v)} \times M$ genotype matrix and $\boldsymbol{b}$ is the $M \times K$ matrix for SNP effects, $\boldsymbol{w} = [w_1, \dots, w_K]^T$ is a $K$-dimensional linear combination weights vector. For simplicity, we assume $\boldsymbol{Y}_1^{(v)}$ is centered, $\boldsymbol{X}_1^{(v)}$ is standardized, and $\boldsymbol{b}$ quantifies standardized SNP effects.

The ridge regression estimator for $\boldsymbol{w}$ is

$$\widehat{\boldsymbol{w}}_{ridge,\lambda} = \underset{\boldsymbol{w}\in\mathbb{R}^K}{\operatorname{argmin}} \|\boldsymbol{Y}_1^{(v)} - \boldsymbol{PRS}^{(v)}\boldsymbol{w}\|_2^2 + \lambda\|\boldsymbol{w}\|_2^2,$$

where $\lambda$ is the shrinkage parameter. It has a closed-form solution:

$$\begin{aligned}\widehat{\boldsymbol{w}}_{ridge,\lambda} &= \left[\boldsymbol{b}^T\boldsymbol{X}_1^{(v)T}\boldsymbol{X}_1^{(v)}\boldsymbol{b} + \lambda\boldsymbol{I}_K\right]^{-1}\boldsymbol{b}^T\boldsymbol{X}_1^{(v)T}\boldsymbol{Y}_1^{(v)} \\ &\approx \left[\frac{N_1^{(v)}}{N^{(ref)}}\boldsymbol{PRS}^{(ref)T}\boldsymbol{PRS}^{(ref)T} + \lambda\boldsymbol{I}_K\right]^{-1}\boldsymbol{b}^T\boldsymbol{X}_1^{(v)T}\boldsymbol{Y}_1^{(v)}.\end{aligned}$$

Here, $\boldsymbol{PRS}^{(ref)} = \boldsymbol{X}^{(ref)}\boldsymbol{b}$, $\boldsymbol{X}^{(ref)}$ is a $N^{(ref)} \times M$ standardized genotype matrix in reference panel. A common practice to obtain the ridge regression estimator is to use two disjoint validation set, one to select the optimal tuning parameter $\lambda$ and the other to estimate $\boldsymbol{w}$ with the selected tuning parameter.

Next, we proposed a summary statistics-based ridge regression for combining multiple PRS. Given the GWAS summary statistics with sample size $N_1$, we subsample GWAS summary statistics for the training set $\boldsymbol{X}_1^{(tr)T}\boldsymbol{Y}^{(tr)}$ with sample size $N_1 - N_1^{(v)}$, and for the validation set $\boldsymbol{X}_1^{(v)T}\boldsymbol{Y}_1^{(v)}$ with sample size $N_1^{(v)}$. We further use to $\boldsymbol{X}_1^{(v)T}\boldsymbol{Y}_1^{(v)}$ to subsample GWAS for two disjoint validation sets: $\boldsymbol{X}_1^{(v1)T}\boldsymbol{Y}_1^{(v1)}$ for validation set 1 with sample size $N_1^{(v_1)}$ and $\boldsymbol{X}_1^{(v_2)T}\boldsymbol{Y}_1^{(v_2)}$ for validation set 2 with sample size $N_1^{(v_2)} = N_1^{(v)} - N_1^{(v_1)}$.

We first apply the PRS method using $\boldsymbol{X}_1^{(tr)T}\boldsymbol{Y}^{(tr)}$ as training data to obatin SNP effects $\boldsymbol{b}$. Next, we obtain grid of the ridge regression estimate $\widehat{w}_{ridge,\lambda}$ using $\boldsymbol{X}_1^{(v1)T}\boldsymbol{Y}_1^{(v1)}$ from validation set 1 and grid of value $\lambda \in \Lambda$. The $\lambda$ with respect to the largest correlation between phenotype and linearly combined PRS in validation set 1 will be used:

$$\hat{\lambda} = \text{argmax}_{\lambda \in \boldsymbol{\Lambda}} \frac{\widehat{\boldsymbol{w}}_{ridge,\lambda}^T \boldsymbol{b}^T \boldsymbol{X}_1^{(v1)T} \boldsymbol{Y}_1^{(v1)} / N_1^{(v_1)}}{\left( \widehat{\boldsymbol{w}}_{ridge,\lambda}^T \boldsymbol{PRS}^{(ref)T} \boldsymbol{PRS}^{(ref)} \widehat{\boldsymbol{w}}_{ridge,\lambda} / N^{(ref)} \right)^{1/2}},$$

where $\boldsymbol{PRS}^{(ref)} = \boldsymbol{X}_1^{(ref)} \boldsymbol{b}$, $\boldsymbol{X}_1^{(ref)}$ is the $N^{(ref)} \times M$ standard genotype matrix in target population reference panel

The final ridge regression-based linear combination weights is estimated in validation set 2 using the formula below:

$$\hat{\boldsymbol{w}}_{ridge,\hat{\lambda}} = \left[ \frac{N_1^{(v_2)}}{N^{(ref)}} \boldsymbol{PRS}^{(ref)T} \boldsymbol{PRS}^{(ref)} + \hat{\lambda} \boldsymbol{I}_K \right]^{-1} \boldsymbol{b}^T \boldsymbol{X}_1^{(v_2)T} \boldsymbol{Y}_1^{(v_2)}.$$

where $\boldsymbol{PRS}^{(ref)} = \boldsymbol{X}^{(ref)} \boldsymbol{b}$, $\boldsymbol{X}^{(ref)}$ is a $N^{(ref)} \times M$ standardized genotype matrix in reference panel.

To avoid overfitting, we recommend using distinct reference panel in summary statistics sampling, PRS model training, ridge regression hyperparameter $\lambda$ selection, and linear combination weights estimation.

##### 3.6 Transforming allele count scale SNP effects into standardized scale

For simplicity, we assume that the genotype matrix is standardized, thus the SNP effects should be on standardized allele scale. Since many PRS method outputs the allele count SNP effects , we use allele frequency from the target population to transform the allele count SNP effects to the standardized effects. The $k$-th column $\boldsymbol{b}_k$ in $M \times K$ SNP effects matrix $\boldsymbol{b} = [\boldsymbol{b}_1, \ldots \boldsymbol{b}_K]$ is calculated by

$$\boldsymbol{b}_k = \frac{\boldsymbol{b}_k^{(\text{allele})}}{\sqrt{2\boldsymbol{f}_1 (1 - \boldsymbol{f}_1)}},$$

where $\boldsymbol{b}_k^{(\text{allele})}$ is the allele count SNP effects for $M$ SNPs in $k$-th population, $\boldsymbol{f}_1$ is the $M$-dimensional target population allele frequency vector. When the in-sample allele frequency is not available, we estimated it from the target population reference panel.

##### 3.7 GWAS summary statistics-based cross-validation

Suppose we divide the full GWAS sample $(\boldsymbol{X}_1, \boldsymbol{Y}_1)$ into a training set $\left(\boldsymbol{X}_1^{(tr)}, \boldsymbol{Y}_1^{(tr)}\right)$ with $N_1 - N_1^{(v)}$ individuals, and a validation set $\left(X_1^{(v)}, Y_1^{(v)}\right)$ with $N_1^{(v)}$ individuals. Given the association z-scores $\left(\frac{\boldsymbol{X}_1^T \boldsymbol{Y}_1}{\sqrt{N_1}}\right)$ from GWAS summary statistics and genotype data from the reference panel, association summary statistics based on training and validation sets can be sampled as:

$$\frac{\boldsymbol{X}_1^{(tr)T}\boldsymbol{Y}_1^{(tr)}}{N_1 - N_1^{(v)}} = \frac{\boldsymbol{X}_1^T\boldsymbol{Y}_1}{N_1} + \left(\frac{N_1^{(v)}}{N_1 - N_1^{(v)}}\right)^{\frac{1}{2}} \frac{\boldsymbol{X}^{(ref)T}}{\sqrt{N^{(ref)}}}\boldsymbol{g}$$

$$\frac{\boldsymbol{X}_1^{(v)T}\boldsymbol{Y}_1^{(v)}}{N_1^{(v)}} = \frac{\boldsymbol{X}_1^T\boldsymbol{Y}_1 - \boldsymbol{X}_1^{(tr)T}\boldsymbol{Y}_1^{(tr)}}{N_1^{(v)}}$$

where $\boldsymbol{X}^{(ref)}$ is a $N^{(ref)} \times M$ standardized genotype matrix from the reference panel for the target population, $N^{(ref)}$ is the sample size of the reference panel, $\boldsymbol{g}$ is a $N^{(ref)}$-dimensional vector with elements drawn from a standard normal distribution.

To perform $P$-folds cross-validation, we first uses the above formula to sample $\boldsymbol{X}_{1,p}^T\boldsymbol{Y}_{1,p}, p = 1, \ldots P-1$ from $P$ independent subset with sample size $\left[\frac{N_1}{P}\right]$ and obtain the GWAS summary statistics from training and validation sets in fold $p$ as:

$$\frac{\boldsymbol{X}_{1,p}^{(tr)T}\boldsymbol{Y}_{1,p}^{(tr)}}{N_1 - \left[\frac{N_1}{P}\right]} = \frac{\boldsymbol{X}_1^T\boldsymbol{Y}_1 - \boldsymbol{X}_{1,p}^T\boldsymbol{Y}_{1,p}}{N_1 - \left[\frac{N_1}{P}\right]}$$

$$\frac{\boldsymbol{X}_{1,p}^{(v)T}\boldsymbol{Y}_{1,p}^{(v)}}{\left[\frac{N_1}{P}\right]} = \frac{\boldsymbol{X}_{1,p}^T\boldsymbol{Y}_{1,p}}{\left[\frac{N_1}{P}\right]},$$

and estimate the linear combination weights in each fold.

### 4 Implementation of other methods

**XPASS** XPASS (Cai et al., 2021) is an empirical Bayes-based PRS framework that leverages genetic correlation for cross-population polygenic prediction. In our paper, XPASS is used to compute heritability and cross-population genetic covariance (correlation), and estimate the SNP posterior effects used to calculate PRS. We used population-matched 1000 Genomes Project data as the reference panel. Five principal components of genotypes in reference panel were used as covariate files as suggested by the software. We estimated the global genetic correlation using genome-wide SNPs. We also created two SNP sets: SNPs inside and outside significant genome regions identified by X-Wing, and computed cross-population genetic correlation using GWAS summary statistics restricted to the two SNP sets separately. Standard errors of genetic parameters (heritability, genetic covariance, and genetic correlation) were estimated using block-wise jackknife method. For PRS construction, we obtained the posterior effects for each population to generate population-specific PRS. Although XPASS did not propose to linearly combine PRS, we applied the linear combination to XPASS-derived PRS for a fair comparison.

**PESCA** As suggested by PESCA paper (Shi et al., 2020), we pruned SNPs such that correlation between SNPs does not exceed 0.95 in the population-matched 1000 Genomes Project data. We used ldetect (Berisa and Pickrell, 2016) to produce LD blocks that are approximately independent in both populations. 14,630 SNPs and 31 independent LD blocks on chromosome 22 were used in simulations; Approximately 500,000 SNPs and 1,368 independent LD blocks were used in GWAS analysis of 31 complex traits. Maximum number of EM iterations to estimate genome-wide prior probability was set as `100` with flag `-max_iter`. Number of independent MCMC chains in estimating posteriors was set as `20` with flag `-max_iter_post`. Number of burn-ins and samples for the MCMC were both set as `5000` by default with flags `-nburn` and `-nsample`, respectively. SNPs with posterior probability larger than 0.95 were identified as shared causal SNPs across populations. To compare findings between X-Wing and PESCA, we extended SNPs identified by PESCA into regions with equal size, such that the aggregated size was the same as that of X-Wing.

**PolyFun-pred** PolyFun-pred (Weissbrod et al., 2022) uses SNP effects estimated from functionally informed fine-mapping to calculate PRS. We downloaded the pre-computed PRS coefficients for PolyFun-pred and used these coefficients to generate PRS. After overlapping with the trait list for the GWAS summary statistics we used, there are 16 traits for East Asians and 11 traits for Americans left.

**PRS-CSx** PRS-CSx (Ruan et al., 2022) is a Bayesian cross-population PRS framework with shared continuous shrinkage across the different populations. We ran PRS-CSx using the default population-matched 1000 Genomes Project data as the reference panel. We used hyperparameter `a=0.5` and `b=0.5` in PRS-CSx, which is exactly the horseshoe prior, for a fair comparison with our method. This choice of the value of `a` and `b` performed almost the same as the default for PRS-CSx (`a=1`, `b=0.5`) given the observation in Table S9 of PRS-CSx paper (Ruan et al., 2022). The global shrinkage parameter was obtained from full Bayesian approach (automatically estimated from data) or model-tuning strategy (value among $\{10^{-6}, 10^{-4}, 10^{-2}, 1\}$ that gives the largest $R^2$ in the validation set will be used).
